# MifRix: An Integrated microbiome framework for predicting generic and disease-specific risks investigating inter-disease diagnostic cross-talks and intra-disease signature variability

**DOI:** 10.64898/2026.08.11.744126

**Authors:** Sourav Goswami, Nalin Arora, Alisha Ansari, Debjit Pramanik, Pavit Singh, Dinesh Palanimuthu, Tarini Shankar Ghosh

**Author notes:** Equally contributing authors (Sourav Goswami, Nalin Arora).

## Abstract

The human gut microbiome is increasingly recognized as a diagnostic indicator across diverse diseases, yet unified frameworks integrating taxonomic and functional features for multi-disease risk assessment, while capturing disease-specific variability in microbiome-alteration signatures, remain limited. Here we present MifRix (Microbiome-inferred Risk-scores with explainability), a two-step ensemble machine-learning framework integrating microbial composition with composition-derived functional signatures to predict generic and disease-specific risk scores across 10 major diseases, coupled with profiling of risk-explainable microbiome features. MifRix was trained using 38,054 gut microbiomes spanning 150 cohorts and 48 nationalities, leveraging taxa abundance and taxa-inferred functional profiles derived from 57,743 functional features mapped across 4,814 species-level taxa. On unseen validation datasets (4,649 microbiomes, 32 cohorts), MifRix outperformed established microbiome health metrics, with disease-specific risk scores achieving strong discrimination (AUC: 0.92-0.99). The explainability module revealed shared microbial signatures across disease pairs, organizing all ten diseases along a continuous gastrointestinal-to-neurological risk gradient, driven not by whole-disease microbiome alteration signatures but by specific, reproducible sub-signatures within each disease. Applied across independent cohorts, MifRix scores further identified population subgroups and individuals at elevated risk for related diseases, flagged precursor/pre-disease states, and tracked therapy-associated response, establishing an interpretable framework for microbiome-based precision diagnostics and disease-risk stratification.

## Introduction

The human microbiome plays a critical role in host physiology and disease susceptibility, with ‘dysbiotic’ microbial signatures consistently associated with almost all major diseases^1–3^. Deviations from a stable microbial configuration, commonly termed dysbiosis, are reproducibly linked to diverse pathological states, including inflammatory bowel disease, colorectal cancer, cardiometabolic disorders, and neurodegenerative diseases^4–9^. As population-scale metagenomic datasets have expanded, machine learning (ML) approaches have increasingly been developed to classify disease states and derive microbiome-based screening tools for predicting individual disease susceptibility^8,10–14^, marking a transition toward predictive, translational applications. Accordingly, multiple microbiome-derived health indices and dysbiosis scores have been proposed to quantify deviation from a healthy microbiome and stratify disease risk^4,7,15–17^, while studies leveraging Explainable Artificial Intelligence (XAI) have sought to delineate interpretable features underlying disease predictions, enhancing model transparency and credibility^18,19^. Although most approaches have focused on taxonomic predictors, function-based wellness prediction tools have recently emerged, reflecting the view that functional capacity offers a more robust diagnostic readout by directly capturing host-microbiome biochemical interactions and their contributions to disease etiology^20–22^. Parallel efforts have characterized microbiome signatures of specific diseases and developed multi-class classifiers capable of assessing multiple disease risks simultaneously^13,23,24^.

However, current gut microbiome-based diagnostic frameworks have several limitations. Most are restricted to a single sequencing platform, either 16S rRNA or whole-genome shotgun metagenomics. Moreover, gut microbiome composition is strongly influenced by geography, industrialization, and lifestyle, leading to substantial population-specific variation^25,26^. Yet, many existing frameworks are trained on population-biased datasets, limiting cross-cohort integration and the robustness and reproducibility of disease signatures. A unified, large-scale diagnostic framework trained on globally diverse microbiome datasets spanning sequencing platforms and populations, while integrating taxonomic and functional features for both generic and disease-specific risk prediction, remains an unmet need.

Furthermore, although prior studies have shown that multiple diseases share partially overlapping microbial signatures, suggestive of conserved dysbiotic axes^13,27^, the extent to which these alterations are shared across disease pairs remains unclear. Diseases vary in mechanistic overlap, with cardiometabolic conditions showing greater similarity than others, a pattern likely shared by host-associated microbiomes. Identifying these shared signatures could reveal cross-disease susceptibilities and therapeutic targets.

Another underexplored aspect is the heterogeneity of microbiome signatures within the same disease. Several cross-cohort studies have shown substantial variation in the performance of microbiome-based disease prediction models across cohorts and populations, influenced by geography, age, and other host-associated factors^28,29^. Thus, disease-associated diagnostic signatures can differ substantially across individuals, with a single disease potentially characterized by multiple distinct patterns of microbiome alteration. However, systematic investigation of this within-disease variability remains largely lacking.

To address platform and population bias, unresolved cross-disease signature sharing, and within-disease heterogeneity, we developed MifRix (Microbiome-inferred Risk-scores with explainability), a two-stage ensemble machine-learning framework that integrates taxonomic composition and taxonomy-derived functional profiles to estimate generic disease (GD) and disease-specific (DS) risk scores from a single microbiome. Trained on gut microbiomes from 48 nationalities across 16S and whole-genome shotgun sequencing platforms, MifRix accurately distinguished healthy from diseased individuals and identified disease-specific patients in an independent validation dataset of 4,692 gut microbiomes from 32 cohorts. MifRix risk profiles further revealed inter-disease relationships consistent with known disease co-occurrence patterns. Explainability analyses identified geographically distinct microbial predictors for the same disease across cohorts, indicating that alternative microbial configurations can underlie similar disease phenotypes. Across nine independent cohorts (n = 1,905 microbiomes), MifRix also captured therapeutic response, personalized treatment outcomes linked to microbiome heterogeneity, longitudinal disease dynamics, and early risk signals from precursor conditions. Together, these findings establish MifRix as a scalable and interpretable framework for microbiome-based disease risk prediction, response stratification, and cross-disease risk profiling.

## Results

### Collation of a global microbiome repository and a comprehensive catalog of microbiome-associated genomes, systematically annotated using multiple functional annotation schemas

The MifRix workflow was designed to jointly leverage microbiome taxonomic abundance and taxa-derived functional representations for robust cross-disease prediction. Its development and validation required a curated global microbiome repository spanning diverse geographic regions, disease groups, and profiling strategies. We therefore compiled 44,533 gut microbiomes from 189 study cohorts across 48 nationalities, including 27,323 controls and 17,210 diseased subjects representing 21 disease categories (**Table S1**). Of these, 18,751 samples (42%) from 96 studies were profiled by whole-genome sequencing (WGS), while 25,782 (58%) from 93 studies were profiled by 16S rRNA amplicon sequencing (16S). The repository captures broad lifestyle diversity, from hunter-gatherer and rural communities to urban, industrialized, mixed, and transitional populations across Africa, Asia, Europe, North and South America, and Oceania.

These 44,533 gut microbiomes were divided into three groups: training dataset comprising 38,054 gut microbiomes from 150 study-cohorts of which 75% were utilized for training the models and 25% for testing; unseen validation datasets comprising 4,649 gut microbiomes from 32 ‘unseen’ study cohorts not considered for training; additional validation dataset consisting of 1,905 gut microbiomes from 9 cohorts not considered for training.

Since microbiome functional profiles were central to our risk-prediction workflow, we constructed a probabilistic association index linking functional annotations to microbiome-associated taxa identified from the taxonomic profiles (**Methods, Text S1**). Briefly, we curated 71,238 high-quality genomes, including metagenome-assembled genomes (MAGs) and reference/isolate genomes, spanning 4,814 species-level taxa (mean 15 genomes per taxon). Using a combinatorial annotation pipeline (**Methods, Text S1**), we generated abundance-based feature vectors comprising 57,743 functional tags across eight annotation categories, including CAZy, BiGG, COGs, EC numbers, InterPro, and KEGG-derived functions (**Fig. 1A**). We then retained the 1,844 species-level taxa detected in our 44,533-microbiome dataset (hereafter, abundance profile; AP), accounting for ≥85% relative abundance in 87% of samples (**Fig. S1**). These taxa were predominantly assigned to *Bacillota* (formerly *Firmicutes*), *Pseudomonadota* (*Proteobacteria*), *Actinomycetota* (*Actinobacteria*), and *Bacteroidota* (*Bacteroides*) (**Fig. 1B**), represented by reference genomes, MAGs, or both. Genome-level annotations were averaged across genomes within each taxon to generate a species-level functional profile matrix.

**Fig. 1.**
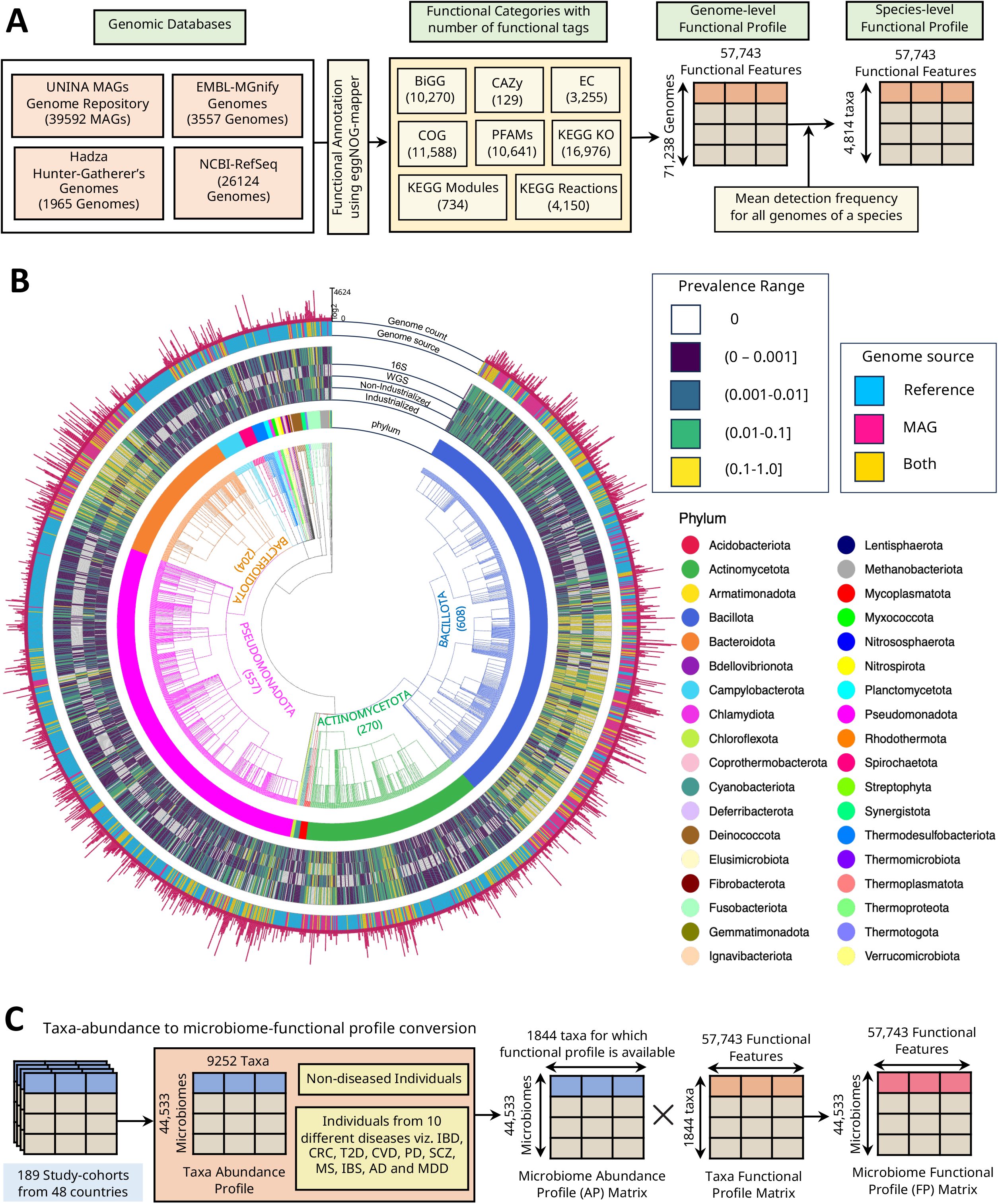
Construction of microbiome-level functional profile by performing a systematic curation of 71,238 high quality genomes across multiple complementary functional categories. A. Schematic description of the protocol utilized for the creation of the species-taxa-level probabilistic functional profile. It provides a consortium of databases utilized for the collection of 71,238 high-quality genomes (encompassing 4,814 unique species-level taxa). These genomes were then annotated with different functional categories (from eight different functional schemas), resulting in total 57,743 functional features or tags detected across at least one genome. The genome-level function abundance matrix (71,238 genomes × 57,743 functional features) was then aggregated using mean at the species-taxa-level to generate mean functional abundance matrix encompassing the abundances of 57,743 functional tags across 4,814 taxa. B. The circular dendrogram-based tree of 1,844 (of the 4,814) taxa was detected at least once in the collection of 44,533 microbiomes curated across the globe, clustered based on their phylogeny and the phylum they belong to. From the innermost ring to the outermost, the circular plot presents the phylum, followed by prevalence of the taxa in the microbiomes from the Industrialized population, prevalence in non-Industrialized population, prevalence in WGS and 16S microbiomes, their genome types (whether a species has genomes as Metagenome-Assembled Genomes (MAGs) or Reference Genomes or both) and finally the count of genomes per species in a log_2_ scale. These 1844 unique taxa belong to 36 different phyla as shown in colour legends. C. This panel shows how the microbiome-level functional profiles have been generated using the cross multiplication between taxonomic abundance profiles of 44,533 microbiomes (from apparently healthy individuals and from individuals encompassing ten different diseases) and the functional profile (consisting of 57,743 different functional tags) of 1844 taxa.

Finally, microbiome-level functional profiles were generated by multiplying the taxonomic abundance matrix (AP; 44,581 microbiomes × 1,844 taxa) with the taxa-to-function abundance matrix (1,844 taxa × 57,743 functional tags), yielding a microbiome-to-function matrix containing the probabilistic abundances of 57,743 functional tags across 44,448 gut microbiomes (hereafter, functional profile; FP) (**Fig. 1C**).

### Devising the MifRix framework

The next step involved the development of the MifRix workflow using 75% of the 38,054 gut microbiomes from 150 study cohorts allocated to the training set (described above), while the remaining 25% were reserved for internal testing. To leverage both microbiome taxonomic composition and taxonomy-derived functional profiles for generic and disease-specific risk prediction, we implemented two complementary variants: MifRix-AP, based on taxonomic abundance, and MifRix-FP, based on taxonomy-derived functional profiles. Both variants employ a two-stage ensemble machine learning framework for risk prediction.

The first stage, referred as the ‘Generic-Disease’ (GD) model, is based on the consistent observations by previous studies that multiple diseases share similar microbiome alterations that can be utilized to ‘diagnose’ a generic state of the gut microbiome negatively associated with health^4–6,27,28^. The GD model predicts a generic-disease risk-score that provides a quantitative measure of how much an individual’s gut microbiome deviates from a health-associated profile and exhibits microbiome signatures shared by multiple diseases. This risk-score is similar to other known measures of microbiome-wellness or dysbiosis^4,7,15,16^. The second phase, referred to as ‘Disease-Specific’ (DS) model, utilizes microbiome alteration patterns (taxonomic for MifRix-AP; functional for MifRix-FP) for a given disease as compared to other diseases to predict a risk-score for that disease. Currently, MifRix incorporates DS models that can predict risk for ten distinct disease categories, namely Colorectal Cancer (CRC), Cardiovascular Diseases (CVD), Inflammatory Bowel Disease/Gut Inflammation (IBD/Gut Inflammation), Irritable Bowel Syndrome (IBS), Parkinson’s Disease (PD), Multiple Sclerosis (MS), Schizophrenia (SCZ), Type 2 Diabetes (T2D), Alzheimer’s Disease (AD) and Major Depressive Disorder (MDD) (refer to **Fig. 2A** and the **Methods** section for detailed selection criteria).

**Fig. 2.**
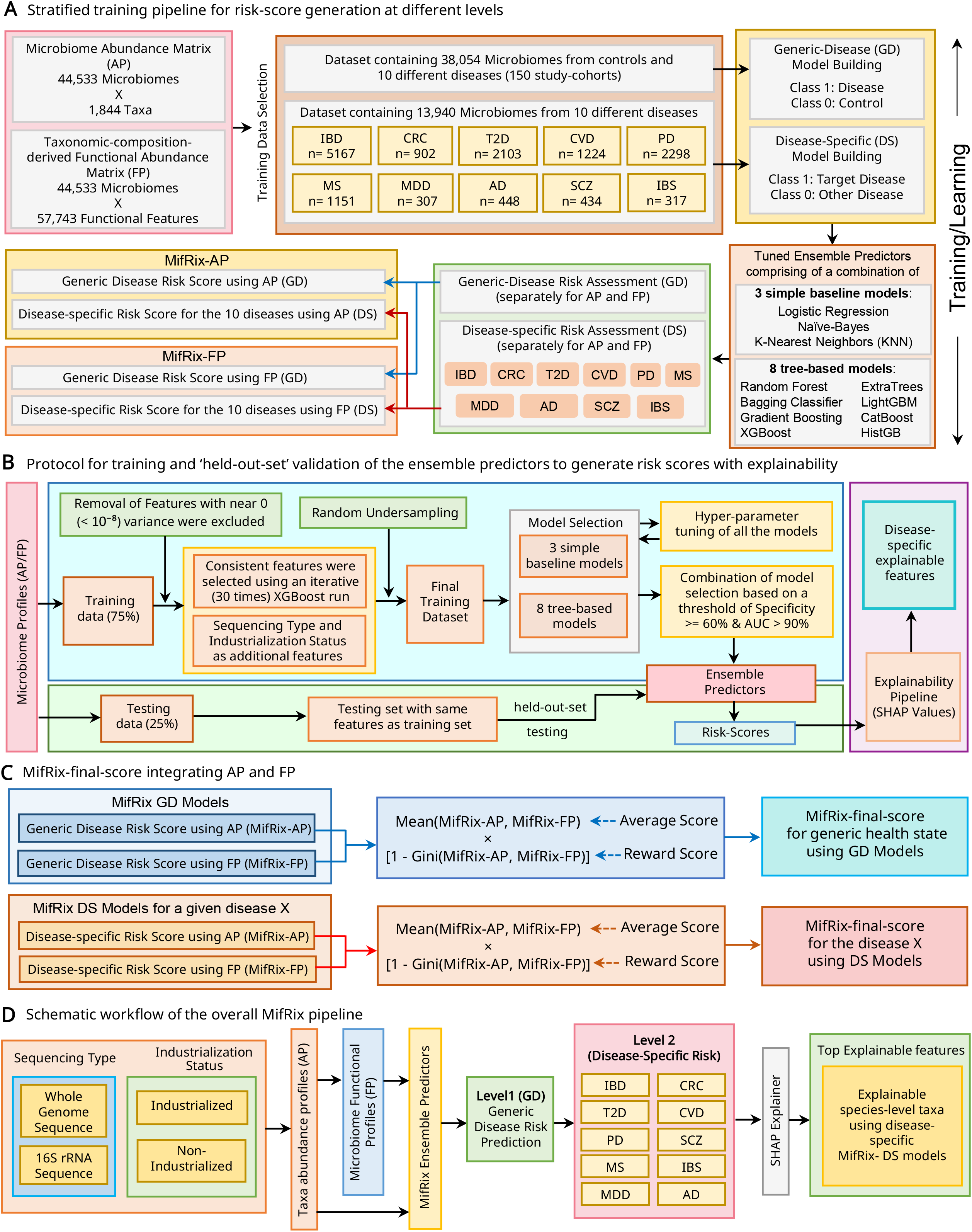
Schematic flow depicting the organization, development, and testing of the MifRix framework. **A.** Overview of the strategy used to build the risk-score generation pipeline, detailing the two input types utilized: the microbiome abundance profile (AP; abundances of 1,814 species-level taxa across 44,533 gut microbiomes) and the taxonomic-composition-derived functional abundance matrix (FP; estimated abundances of 57,743 functional tags across the same microbiomes). Of these, 38,054 microbiomes across 150 study-cohorts (including >13,000 microbiomes spanning the 10 diseases) were selected for the training step. For both AP and FP, 11 distinct model sets were trained, one generic-disease model (GD) predicting overall disease risk and ten disease-specific models (DS) predicting independent risk for each of the 10 diseases, as in the schematic. **B.** Detailed schematic of the training protocol. The 38,054 microbiomes across 150 cohorts were divided into a 75% training subset (used to train the models) and a 25% held-out testing subset. For each of the 22 models, features with variance <10 were removed, followed by iterative XGBoost-based feature selection. Sequencing type and industrialization status were additionally included as features to account for sequencing platform and population-level heterogeneity across study-cohorts, followed by random under-sampling to address class imbalance across datasets. An ensemble model was then built by training 11 individual models, with hyperparameter tuning used to select the best-performing model combination, balancing specificity and recall to yield the final generic and disease-specific ensemble predictors for both AP and FP. **C.** Schematic workflow for developing the combinatorial MifRix-final-score, which integrates the MifRix-AP and MifRix-FP frameworks into a single unified measure, computed as the mean of the AP- and FP-derived risk scores multiplied by (1 - Gini), thereby down-weighting microbiomes showing greater inequality between the two risk-score variants. **D.** Describes the overall MifRix pipeline. A new microbiome (irrespective of 16S/WGS or industrialized/non-industrialized origin) receives risk predictions at both the generic-disease (GD) and disease-specific (DS) levels, using both taxa abundance profiles (AP) and microbiome-functional profiles (FP), with the top explainable features driving each prediction also identified. This full framework was applied to assess MifRix performance across the 32 unseen validation datasets.

Overall, MifRix comprises 22 risk-prediction models: 11 taxonomy-based (MifRix-AP) and 11 taxonomy-derived functional (MifRix-FP) models. Each set includes one generic disease (GD) model and ten disease-specific (DS) models, enabling simultaneous estimation of global and disease-specific risk from microbiome taxonomic composition and functional potential.

To accommodate the complex, non-linear interactions underlying gut microbiome structure and host associations, we adopted an ensemble machine-learning strategy rather than relying on a single algorithm. Each of the 22 models integrated 11 classifiers through a voting-based ensemble, with the optimal algorithm combination selected to maximize predictive performance and robustness (**Fig. 2A**; **Methods**; **Text S2**). The ensemble combined three interpretable baseline methods: Logistic Regression, Naïve Bayes, and K-Nearest Neighbors (KNN) with eight tree-based bagging and boosting algorithms: Random Forest, Extra Trees, Gradient Boosting, HistGradientBoosting, XGBoost, LightGBM, Bagging Classifier, and CatBoost. Following hyperparameter optimization, predictions from individual classifiers were combined using an ensemble predictor that selected the optimal model combination for each risk estimate (**Fig. 2B**). GD and DS models were trained on 75% of the 38,054 microbiomes from the training cohorts and evaluated on the remaining 25% (**Methods**, **Text S2**; **Table S2**).

Notably, MifRix explicitly incorporated the sequencing strategy (16S or WGS) and industrialization status (industrialized or non-industrialized) of each microbiome as model features, allowing it to account for major sources of cross-cohort variation while retaining microbiome-derived disease signals. This design enabled a unified risk-prediction framework spanning both 16S rRNA and whole-genome shotgun datasets and cohorts differing in population industrialization (**Fig. 2B**). We further incorporated a SHAP-based explainability module to quantify feature contributions to model predictions. Finally, for each disease model, we computed a MifRix-final score by averaging the corresponding MifRix-AP and MifRix-FP risk scores and multiplying the result by (1 - Gini), thereby down-weighting predictions with greater disagreement between the two modalities (**Fig. 2C**). The overall MifRix workflow, integrating sequencing type and industrialization status with taxonomic and functional profiles, followed by ensemble-based generic and disease-specific risk prediction and SHAP-based identification of top explanatory features, is summarized in **Fig. 2D**.

### MifRix-GD models outperform state-of-the-art microbiome health indices and MifRix-DS models show strong disease discrimination

We first evaluated the generic disease (GD) predictors, MifRix-AP and MifRix-FP, using the 25% held-out subset of the training cohorts. Both achieved excellent discrimination, with AUCs of 0.94 and 0.95, respectively (**Fig. S2A-B**). Disease-specific (DS) models similarly performed well, with AUCs ranging from 0.93-0.99 for MifRix-AP and 0.92-0.98 for MifRix- FP (**Fig. S2C-D**; **Table S3**). Direct comparison of AP- and FP-based DS models showed comparable performance for distinguishing controls from disease and for discriminating among disease categories within the training cohorts (**Fig. S3**). Collectively, the consistently high AUCs (>0.90) across generic and disease-specific models demonstrate the robustness of MifRix as a microbiome-based disease screening framework.

We next evaluated MifRix on 32 independent study cohorts comprising 4,649 gut microbiomes across 18 diseases, including eight disease classes absent from model training-Acute Diarrhoea, Polyps, Hypertension, COVID-19, Depression, Congenital Chloride Diarrhoea (CLD), Behcet’s Disease (BD), and Postpartum Depressive Disorder (PDD) to assess the generic disease (GD) models beyond the training disease space. Both MifRix-AP and MifRix-FP performed robustly, achieving ≥90% accuracy in eight and nine cohorts, respectively, and ≥80% accuracy in 14 and 15 cohorts, respectively (**Fig. S4A**; **Table S4**). MifRix-FP achieved a modestly higher weighted mean accuracy than MifRix-AP (78% vs. 75%), whereas the MifRix-final score showed intermediate performance by integrating predictions from both models. Classification accuracy was calculated by predicting each microbiome as diseased (GD) when the corresponding GD risk-score was > 0.5, and comparing these predictions against the true disease labels.

We then benchmarked the two MifRix variants and the final-score against established microbiome-based health indices, including the Dysbiosis-Score^15^, Gut Microbiome Health Index (GMHI)^16^, Gut Microbiome Wellness Index 2 (GMWI-2)^7^ and the HACK-Top17-Score^4^ (previously developed by us), across 28 completely unseen cohorts with matched disease and control microbiomes (**Text S3**). Using the Mann-Whitney U test to assess how frequently each index showed significant control-versus-disease differences in the expected direction (higher in disease: Dysbiosis-Score, MifRix-FP, MifRix-AP, MifRix-CS; higher in controls: HACK-Top-17, GMHI, GMWI-2), the GD risk scores of both MifRix variants and the final-score outperformed all other indices, showing significantly higher values in diseased subgroups (p ≤ 0.05) in 16 of 28 cohorts (57%), compared to 13 of 28 (46%) for the Dysbiosis-Score, the next-best performer (**Fig. 3A**).

**Fig. 3.**
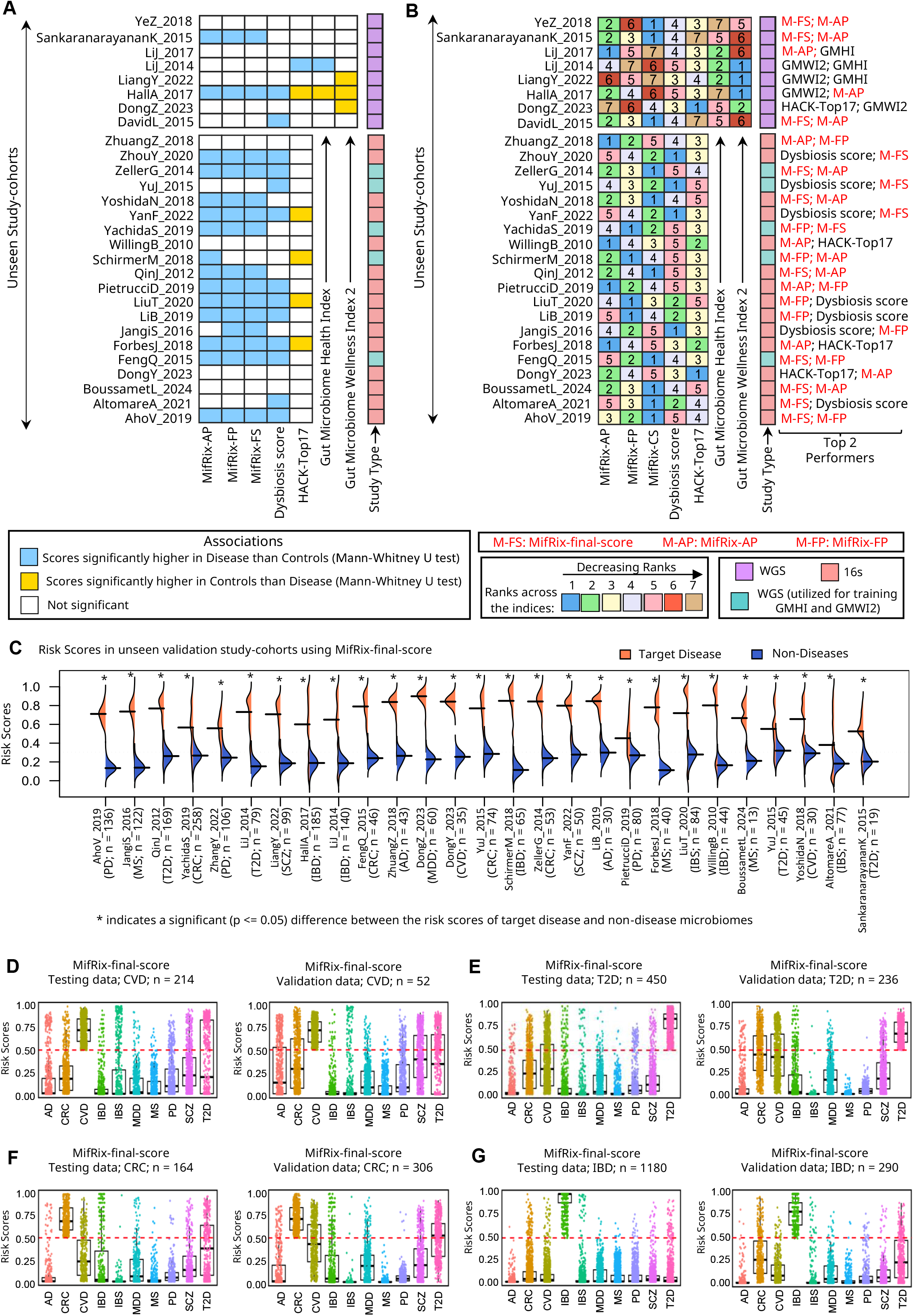
Performance evaluation of the MifRix framework and observation of inter-disease crosstalk. **A-B.** Superior performance of Generic Disease (GD) risk-scores (MifRix-final-score, MifRix-AP, MifRix-FP) in differentiating diseased from non-diseased states compared to state-of-the-art wellness indices. **A.** Heatmap comparing MifRix GD risk-score variants against established indices (Dysbiosis-Score, GMHI, GMWI-2, HACK-Top17-Score) across 28 unseen validation cohorts with matched controls and disease samples. Mann-Whitney tests compared each index between controls and disease; sky-coloured cells indicate significantly higher disease values (p ≤ 0.1), golden cells indicate significantly higher control values (p ≤ 0.1). The predominance of sky-coloured cells shows that MifRix-AP, MifRix-FP, and MifRix-final-score together consistently captured disease-versus-control differences, with the Dysbiosis-Score performing next best. **B.** Comparative SHAP-based ranking of the same indices across the same 28 cohorts, using LightGBM classifiers trained on all indices per cohort; median absolute SHAP values ranked each index’s relative contribution. The ’Study Type’ strip indicates 16S (peach) or WGS (violet) profiling, and whether a cohort was used to train GMHI/GMWI2 (green); top-2 performers per cohort are labelled. MifRix variants consistently ranked among the strongest predictors, with MifRix-final-score ranking first in 10/28 and top-two in 19/28 unseen validation cohorts, including cohorts representing diseases excluded from GD model training. **C.** Split bean plots showing MifRix disease-specific (DS) model performance across 27 unseen validation cohorts spanning the 10 diseases. Left (orange) shows target-disease MifRix-final-score distributions; right (blue) shows mean scores for the nine non-target diseases. Asterisks indicate significant differences (p ≤ 0.05, Wilcoxon rank-sum test). The same distributions for the AP and FP variants are in Figs. S5D and S5E, respectively**. D-G.** Inter-disease crosstalk. Distribution of MifRix-final-score-derived risk-scores across all ten diseases for microbiomes predicted high-risk (score >0.5) for CVD (**D**), T2D (**E)**, CRC (**F**), and IBD (**G**), shown as box plots with jittered points (each point = one microbiome) for testing (left) and validation (right) cohorts. While target-disease risk-scores were consistently highest, specific non-target diseases also showed elevated scores (e.g., SCZ/T2D for CVD; CVD for T2D; T2D/IBD for CRC).

To further benchmark performance, we trained LightGBM (LGBM) classifiers to distinguish control from disease microbiomes in each cohort using all indices as predictors, and ranked their contributions by SHAP analysis (**Methods**; **Text S3**; **Fig. 3B**; **Table S5**). In 25 of 28 cohorts (∼90%), at least one MifRix variant ranked among the top two predictors based on mean SHAP values. The MifRix-final score ranked first in 10 cohorts and among the top two in 19, including four of five cohorts representing disease classes absent from training (Behcet’s disease, Hypertension, Polyps, and Acute Diarrhoea; **Fig. 3B**). These findings highlight the strong generalizability of MifRix for distinguishing healthy and disease-associated microbiome states, including diseases unseen during training.

We next evaluated the disease-specific (DS) models across the 10 diseases implemented in MifRix using 3,675 microbiomes from the training-cohort test set and 2,182 independent microbiomes spanning 27 study-disease combinations from 25 validation cohorts (**Tables S1**, **S4**). Across both datasets, target disease risk scores were consistently higher than the mean risk scores for non-target diseases (Wilcoxon rank-sum test, *P* ≤ 0.05; **Figs. 3C**, **S5A-E**), demonstrating robust disease-specific risk discrimination, including in independent validation cohorts. MifRix-FP showed marginally stronger disease-specific discrimination than MifRix-AP across the validation cohorts (**Fig. S4B**).

### Disease-specific microbiome risk scores reveal inter-disease crosstalk and shared disease susceptibility

Rather than treating diseases as mutually exclusive classes, MifRix models the risk of each disease independently, enabling simultaneous profiling of disease-specific (DS) risk across all ten conditions. As many diseases share biological mechanisms bidirectionally linked to the gut microbiome, this framework offers an opportunity to detect microbiome-associated inter-disease crosstalk that a single-label classifier would obscure.

To test this, we examined the distribution of DS risk-scores across all ten diseases (comparing MifRix-final-scores first), asking whether patients scoring highest for their own disease also showed elevated scores for other conditions (**Table S6**). This proved true for several diseases, revealing microbiome signatures linking specific disease pairs (**Figs. 3D-G**, **4A-F**).

#### Cardiometabolic and gastrointestinal crosstalk

CVD patient microbiomes scored highest for CVD but also disproportionately high for T2D, and the reciprocal pattern held for T2D patients (**Figs. 3D-E**), consistent across testing and validation cohorts. This reciprocal pattern reflects the well-established cardiometabolic continuum, in which shared, mutually reinforcing disruptions to inflammatory and metabolic pathways link the two conditions^30,31^.

A similar pattern emerged among CRC, IBD, and T2D. CRC microbiomes scored elevated for IBD and T2D, while IBD and T2D microbiomes scored disproportionately high for CRC, second only to their own target-disease scores (**Fig. 3E-G**), reproduced across all sub-cohorts. This corroborates prior work linking insulin resistance, chronic inflammation, oxidative stress, and microbiome-mediated metabolic dysregulation to colorectal carcinogenesis^32–34^.

#### Neurological cross-disease links

Extending this analysis to neurological disorders revealed further associations, both with gastrointestinal/metabolic conditions and among neurological diseases themselves (**Fig. 4A-F**). AD microbiomes showed elevated risk-scores for SCZ, PD, and CVD, reciprocally (**Figs. 3D, 4A-C**), supporting reports linking neurodegeneration, vascular dysfunction, psychiatric disorders, lipid dysregulation, and chronic inflammation^35–40^. PD and MS, both linked to neuroinflammation and blood-brain-barrier disruption, showed reciprocal associations, as did PD-SCZ, together corroborating shared neuroinflammatory, dopaminergic, and immune-dysregulation mechanisms^41^(**Fig. 4D**). MDD microbiomes showed relatively elevated risk-scores for AD and SCZ (Fig. 4E). Notably, IBS, spanning both neurological and gastrointestinal/metabolic aetiology, showed elevated scores for PD, SCZ, and also metabolic disorders, consistent with the gut-brain axis and reports of gastrointestinal dysfunction preceding Parkinsonian manifestations^42^ (**Fig. 4F**).

**Fig. 4.**
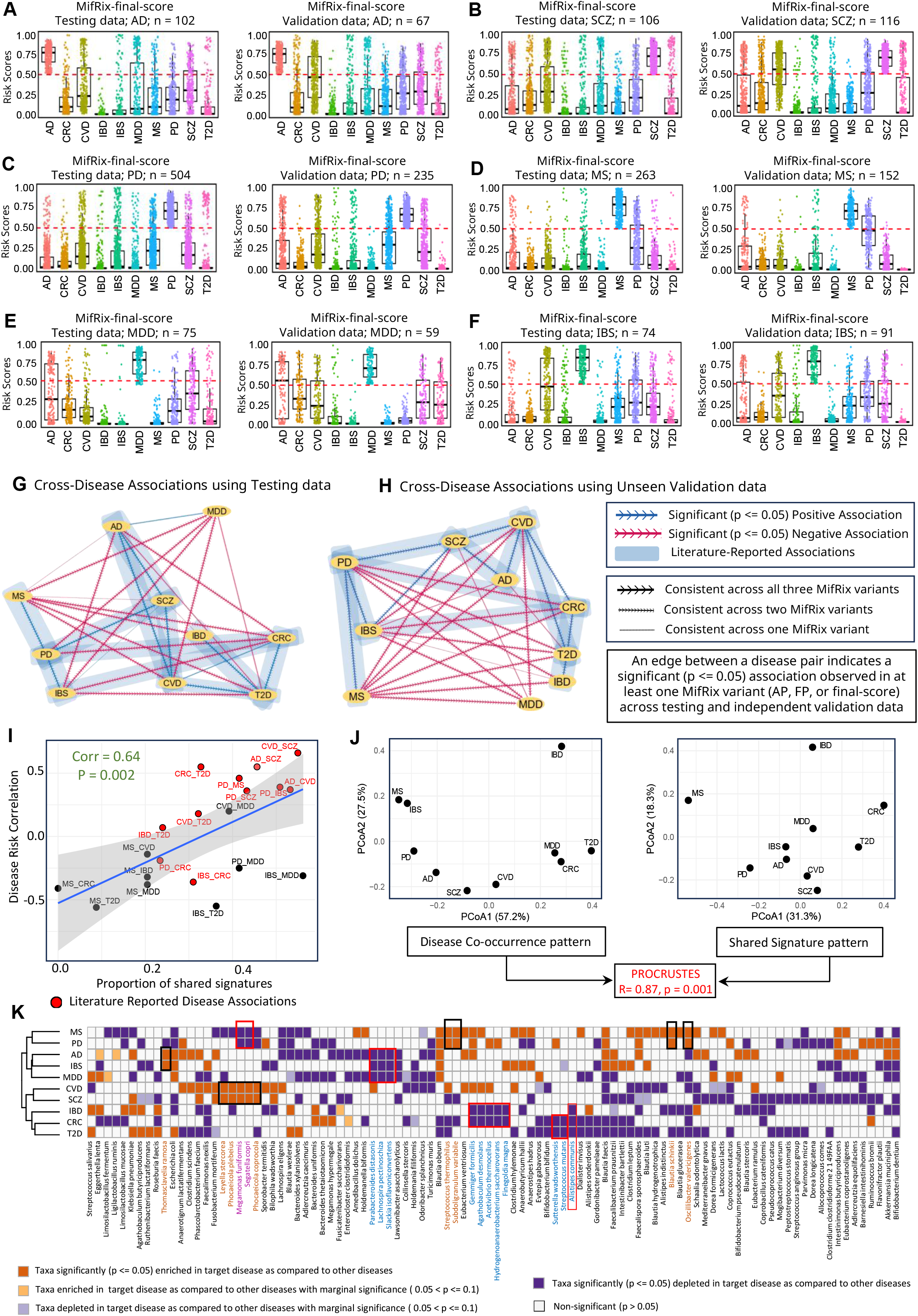
MifRix framework enables identification of inter-disease crosstalk and shared disease susceptibility patterns. **A-F.** Distribution of MifRix-final-scores across all ten diseases for patient microbiomes predicted as high-risk (MifRix-final-score >0.5) for their target diseases where the target diseases are: AD (**A**), SCZ (**B**), PD (**C**), MS (**D**), MDD (**E**), and IBS (**F**), shown for both testing (left) and validation (right) cohorts as box plots with jittered points (each point = one microbiome). Corresponding distributions for CVD/T2D/CRC/IBD appear in Fig. 3D**-G**; and for AP/FP variants in **Figs. S6-S7**. **G-H.** Association networks for testing (G) and validation (H) datasets, integrating MifRix-AP, MifRix-FP, and final-score enrichment results, retaining only associations with consistent directionality across all three. Edge weights reflect significance (p ≤ 0.05) consistency across variants (±1: all three; ±0.7: two; ±0.3: one); positive/negative signs denote enrichment/depletion (blue/red edges). Associations concordant across both networks and all three variants were deemed robust; those additionally literature-supported are marked with blue bands (**Table S13**). **I.** Correlation between shared signature proportion and MifRix-AP disease risk correlations (literature-supported pairs in red); regression (grey, 95% CI) shows significant positive correlation (Pearson’s r = 0.64, p = 0.002). **J.** Procrustes analysis comparing disease-co-occurrence and microbiome-signature ordinations shows strong concordance (r = 0.87, p = 0.001), indicating signature sharing reflects known co-occurrence. **K.** Heatmap of differential taxa across the ten diseases (top 50 species/disease by positive SHAP value), coloured by direction/significance of differential abundance versus all other diseases: dark orange, significant enrichment (Hedges’ g > 0, p ≤ 0.05); light orange, marginal enrichment (Hedges’ g > 0, 0.05 < p ≤ 0.1); dark purple, significant depletion (Hedges’ g < 0, p ≤ 0.05); light purple, marginal depletion (Hedges’ g < 0, 0.05 < p ≤ 0.1); white, non-significant (p > 0.05). Overall, 20 disease-pairs showed significant positive or negative MifRix risk-score relationships, 11 with prior literature support; among the 10 significantly positive pairs, nine had literature-reported co-occurrence.

#### Formal enrichment analysis

To systematically test these relationships, for each target disease we identified microbiomes from the nine other groups that nonetheless scored above 0.5 for the target disease, then used Fisher’s exact test to assess enrichment or depletion relative to expected frequencies (**Methods**). This was performed for all three MifRix variants (final-score, AP, FP; **Tables S7-S12**; **Figs. S6-S7**) across testing and validation cohorts, six tests per disease. Twenty disease-pairs showed significant, consistently directional associations across all six tests (**Fig. 4G-H**), of which 11 (55%) had prior literature support for the same direction of co-occurrence or divergence (**Table S13**).

Reproducible negative associations also emerged. CRC showed a significant inverse relationship with PD, corroborating earlier epidemiological reports^43,44^, and with IBS, suggesting distinct microbial ecosystems and divergent inflammatory or metabolic architectures^43–46^. Together, these findings demonstrate that microbiome-derived risk profiling robustly captures inter-disease crosstalk across diverse disorders, pointing to shared pathophysiological mechanisms across specific disease-pairs. We next investigated the specific taxa mediating these relationships.

### Shared microbial signatures underlie disease co-occurrence patterns

We next assessed whether shared microbial signatures between diseases underlie the pairwise similarity of disease-specific risk-scores. MifRix explains each prediction using SHAP values, which quantify each feature’s contribution to a given DS risk-score. For all downstream analyses, we focused on MifRix final-risk-scores, which captured overall AP/FP association patterns, and on taxonomic features, that were fewer in number (n = 1,844 vs. n = 57,743 functional features) and better characterized than functional features, thereby making shared-feature investigation across diseases more tractable.

For each disease, we ranked features by SHAP score and determined alteration directionality by comparing abundances between target-disease and other samples, yielding disease-specific microbiome signatures (**Methods**, **Fig. S8**), from which we derived a microbial signature similarity score for each disease pair based on overlap and directionality (**Methods**). Across twenty disease pairs with consistent, significant associations (**Fig. 4G-H**), signature similarity correlated strongly with DS risk-score-based correlations (Spearman’s ρ = 0.64, p = 0.002; **Fig. 4I**). Strikingly, 9 of the 10 disease-pairs showing significant positive DS-risk-score correlations were independently corroborated by prior reports of clinical co-occurrence (**Table S13**).

Procrustes analysis of PCoA ordinations confirmed significant concordance between disease-relationship spaces defined by shared signatures and by risk-score co-occurrence (R = 0.87, p = 0.001; **Fig. 4J**), indicating that shared microbial alterations shape the interconnected landscape of human disease.

At finer taxonomic resolution, these signatures revealed biologically meaningful disease groupings (**Fig. 4K**). MS and PD clustered together, sharing enrichment of *Oscillibacter valericigenes*, *Blautia schinkii*, *Streptococcus thermophilus*, and *Subdoligranulum variabile*, alongside depletion of *Megamonas funiformis* and *Segatella copri*, corroborated by multiple prior studies^47–56^. Notably, *S. copri* (formerly *Prevotella copri*) depletion has previously been linked to Lewy body formation and alpha-synuclein aggregation, a hallmark of PD pathogenesis^47^.

A second group comprised AD, MDD, and IBS, a disorder with both gastrointestinal and neurological aetiology, consistent with the gut-brain axis, sharing depletion of *Parabacteroides distasonis*, *Lachnospira pectinoschiza*, and *Slackia isoflavoniconvertens*, alongside enrichment of *Thomasclavelia ramosa* in AD and IBS. *Parabacteroides* depletion is consistently reported across all three conditions in meta-analyses, cohort studies, and preclinical models^57–59^; notably, *Parabacteroides* abundance rises in IBS patients responding to FMT, inversely tracking symptom severity^59^. *Lachnospira* depletion shows a similarly consistent pattern across all three disorders^60,61^.

A third grouping linked SCZ with cardiovascular disease (CVD), characterized by enrichment of *Leyella stercorea*, *Phocaeicola* (*P. plebeius*/*coprocola*), *Megamonas funiformis*, and *S. copri*^62–70^. *S. copri* and *L. stercorea* (both earlier *Prevotella* spp., family Prevotellaceae) have previously been shown enriched in both diseases^71–73^, with *S. copri* causally implicated via Trimethylamine (a TMAO precursor linked to CVD progression) and via acetate production and serotonin dysregulation linked to schizophrenia-like behaviour^72,73^. The final group, IBD, CRC, and T2D, clustered together, reflecting shared inflammatory and cardio-metabolic signatures (**Fig. 4K**), characterized by depletion of *Alistipes communis*, with IBD-CRC additionally marked by depletion of butyrate-producing taxa including *Gemmiger formicilis*, *Agathobaculum desmolans*, *Acetivibrio thermocellus*, *Hydrogenoanaerobacterium saccharivorans*, and *Finegoldia magna*. The CRC-T2D relationship was further distinguished by depletion of *Sutter Ella wadsworthensis* and *Streptococcus mutans*, illustrating both conserved and pair-specific signatures underlying disease co-occurrence.

Together, these findings reveal microbiome alteration signatures shared across distinct, often comorbid disease-pairs, with many individual taxonomic signatures independently corroborated by isolated, disease-specific studies, providing convergent support. Beyond these pairwise relationships, this analysis uncovers a more fundamental principle: individual taxa are not uniformly “beneficial” or “harmful,” but exert disease-specific, context-dependent effects. *S. copri* exemplifies this duality, associated with elevated risk in CVD and SCZ yet reduced risk in PD, suggesting a single microbe can influence divergent disease processes through distinct mechanistic pathways depending on host and disease context, a taxon-level pleiotropy that MifRix’s disease-specific architecture is uniquely positioned to capture.

### Reproducible diagnostic sub-signatures for different diseases confine cross-disease links to specific patient subsets

A key feature of MifRix’s SHAP module is that it computes SHAP values at the individual-sample level, revealing each sample’s own top predictive features rather than a single disease-average signature. This lets us ask whether a disease’s microbiome presents as one shared signature across patients or breaks into distinct sub-signatures driven by different features in different samples, a hypothesis motivated by prior evidence that microbiome classifiers often generalize poorly across cohorts and sub-populations due to host-specific factors^74,75^. This same resolution lets us revisit the inter-disease crosstalk identified earlier at finer detail: we previously showed each disease is characterized by multiple diagnostic signatures variably linked to different comorbid diseases; here we ask whether all of a disease’s signatures carry the same comorbidity risk, or whether each point to a different one.

For each disease, SHAP profiles from correctly predicted (risk-score > 0.5) testing-cohort samples were clustered via k-means to identify distinct microbial sub-signatures, with cluster number chosen for best separation and visualized via PCA; reproducibility in the validation-cohorts was confirmed by projecting validation-cohort SHAP values into the same PCA space and checking cluster mapping. For each sub-signature, the top 20 species driving predictions (by median SHAP value) were identified and their abundance compared to healthy controls and other-disease samples, with consistent, significant directional shifts scored as enriched or depleted, independently in both cohorts (**Methods**, **Figs. S9-S13**).

The number of distinct sub-signatures varied across the 10 diseases (detailed in **Text S2**; key findings summarized here). IBD showed four reproducible sub-signatures, consistently distinct against both controls and other diseases (**Fig. S10A**), of which only two (IBD-signature-1, -2) showed elevated risk-scores for both CRC and T2D. CRC and T2D each showed three similarly reproducible sub-signatures. While CRC-signature-1 was elevated specifically for T2D, CRC-signature-3 specifically for CVD. For T2D, only T2D-signature-1 (of three) showed elevated risk for both CRC and CVD (**Figs. 5**, **S10**). Thus, while IBD, CRC, and T2D showed strong cross-disease risk as a group (**Fig. 4**; consistent with prior literature), this granular analysis revealed that only specific sub-signatures within each disease drive these associations (**Fig. 5**).

**Fig. 5.**
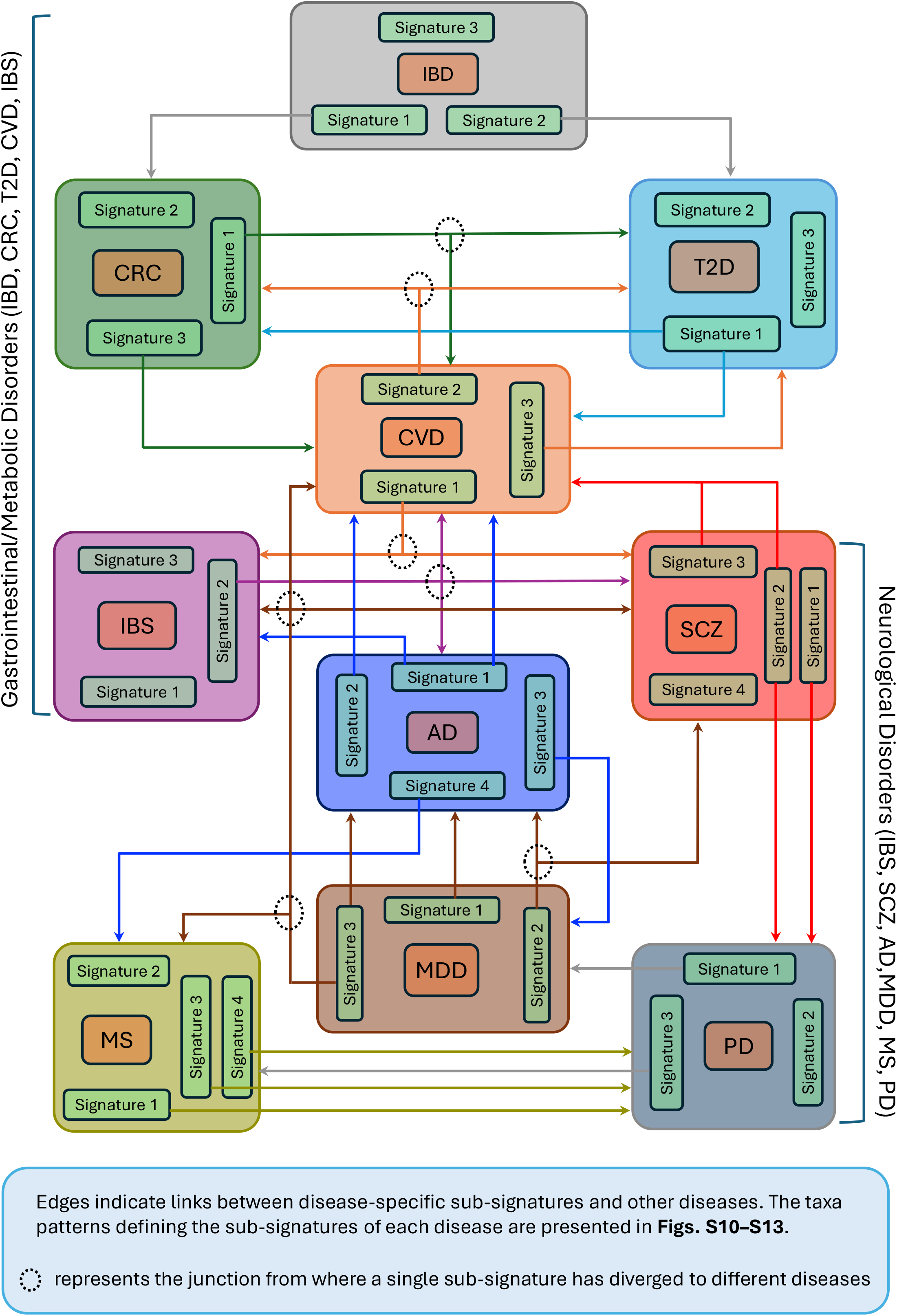
Reproducible diagnostic sub-signatures for different diseases confine cross-disease links to specific patient subsets. Each disease is represented by a box, with its constituent microbiome alteration sub-signatures shown as smaller green boxes nested within it (detailed in **Text S4** and **Figs. S10-S13**). Diseases are arranged according to their inter-disease positive association patterns, with edges drawn from a given sub-signature to the specific diseases for which microbiomes carrying that signature show elevated risk. Black dotted circles mark junctions where a single sub-signature is linked to multiple diseases. This connectivity arranges the 10 diseases along a disease spectrum: gastrointestinal/metabolic disorders (IBD, CRC, T2D) at one end, followed by CVD and IBS (the latter bridging gastrointestinal and neurological manifestations) and neurological disorders (SCZ, AD, MDD, PD, MS) at the opposite end.

CVD had previously shown links to gastrointestinal/metabolic disorders (CRC, T2D) and neurological disorders (SCZ, IBS; **Fig. 4**). Granular analysis identified four CVD sub-signatures. CVD-signature-2 was enriched for CRC and T2D risk; CVD-signature-3 for T2D alone; and CVD-signature-1 for both SCZ and IBS. Reciprocally, only IBS-signature-1 (of three) showed this specificity for CVD, AD, and SCZ (**Figs. 5**, **S11A-B**). For SCZ, two of four sub-signatures (SCZ-signature-2, -3) drove the strongest CVD-risk enrichment, while SCZ-signature-1 instead associated with PD (**Figs. 5**, **S11C**).

At the other end of the spectrum, four neurological disorders, AD, MDD, MS, and PD, again showed cross-disease associations driven by specific, not all, sub-signatures. AD comprised four sub-signatures, AD-signature-1 linked to CVD and IBS, AD-signature-2 to CVD alone, and AD-signature-3/-4 to MDD and MS, respectively (**Figs. 5**, **S12A**). A similar pattern held among PD, MS, and MDD (**Figs. 5**, **S12B**, **S13**).

Together, these findings show that inter-disease associations based on shared microbiome signatures are not random, but track previously reported co-occurrence and comorbidity rates. The 10 diseases could be arranged along a spectrum, gastrointestinal/metabolic disorders (IBD, CRC, T2D), followed by CVD, then IBS (spanning both symptom types), and finally neurological disorders (SCZ > AD > MDD > PD > MS). Critically, cross-disease risk was not uniform across patients with a given disease, but concentrated in those carrying specific microbiome sub-signatures underpinned by distinct taxonomic alterations (**Text S4**).

### MifRix risk-scores flag at-risk subgroups, individuals in precursor disease-states, and enable profiling/measurement of therapeutic response

Our next objective was to probe the translational applicability of MifRix (details in **Text S5**). We first examined its robustness in identifying at-risk sub-populations, using a previously published multi-ethnic cohort of 214 Malaysians (Chinese, Malay, Indian, Jakun) ^76^, in which Indians had the highest reported diabetes burden. MifRix precisely recapitulated this. T2D risk-scores differed significantly across ethnic groups, with Indians showing the highest T2D DS risk-scores (**Fig. 6A**).

**Fig. 6.**
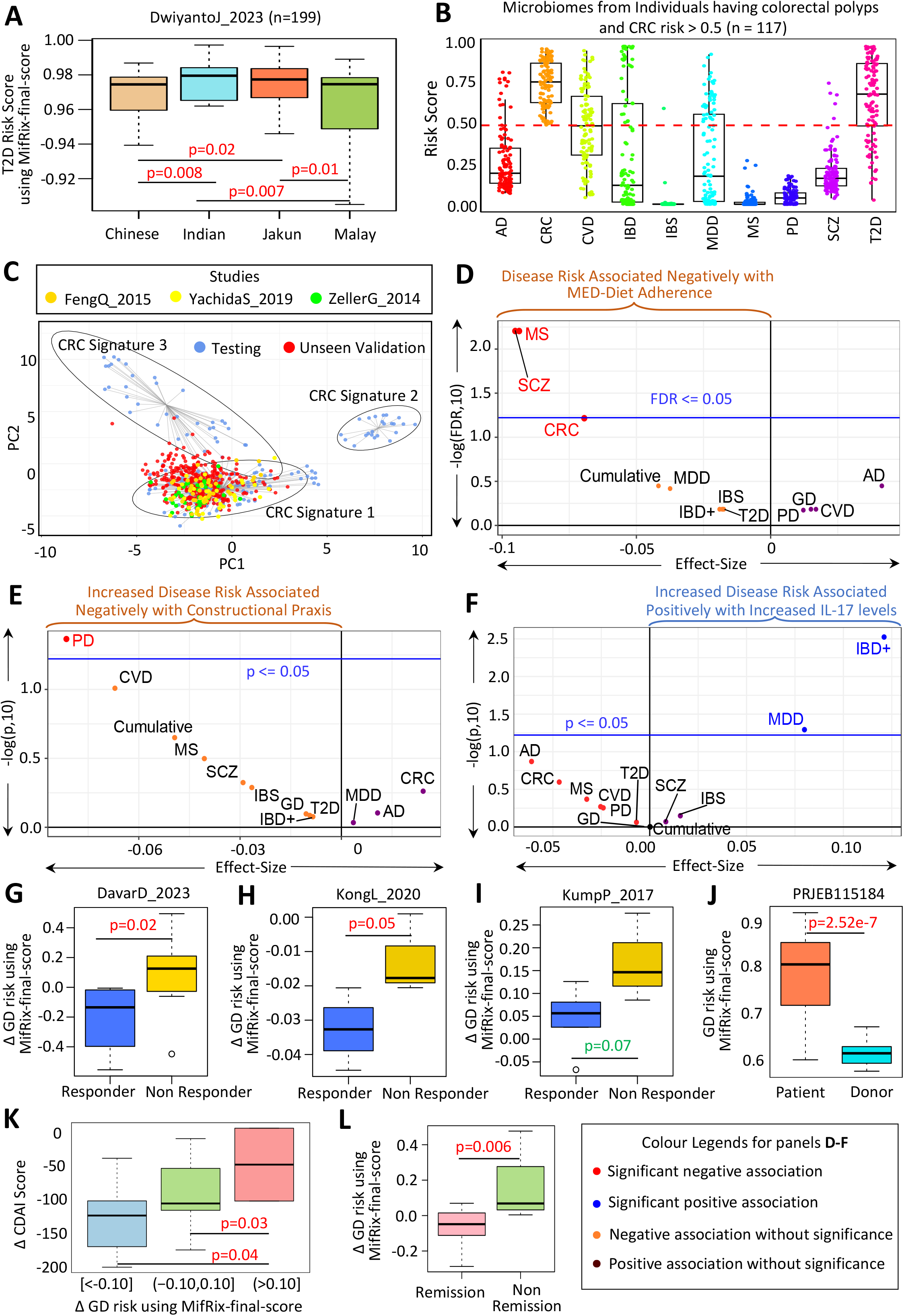
MifRix-derived risk-scores identify at-risk subgroups, individuals in precursor disease-states, and enable profiling of therapeutic response. **A.** T2D-specific MifRix-final-scores across four population groups (Chinese, Indian, Jakun, Malay) in the Dwiyanto_2023 cohort (n=199). Indians showed significantly higher scores, recapitulating the original study’s finding of highest T2D burden in Indians. **B.** MifRix-final-scores for all 10 diseases in colorectal polyp patients (n=117; FengQ_2015, YachidaS_2019, ZellerG_2014), showing expected elevated CRC risk. **C.** Projecting polyp-microbiome based on the SHAP values onto the CRC signature PCA space (different studies’ microbiomes in different colours; **Fig. S10B**) shows most cluster with CRC-signature-1, predominant in both training (blue) and validation (red) CRC microbiomes, consistent with the elevated CVD/T2D risk also seen in this sub-signature. **D–F**. Results from the NU-AGE cohort (GhoshT_2020) evaluating MifRix for therapeutic-efficacy assessment. **D.** Disease-specific risk-scores versus Mediterranean Diet (MED-diet) adherence (x-axis: Pearson correlation; y-axis: -log FDR); adherence was negatively associated with risk for 7/10 diseases, significantly for CRC, MS, SCZ. **E.** Change in Constructional Praxis scores versus change in disease-risk; increases linked to significantly decreased Parkinson’s risk. **F.** Change in IL-17 (pro-inflammatory cytokine) versus change in disease-risk; increases significantly positively associated with IBD/gut-inflammation risk. For **E–F**, axes show effect-size (Pearson correlation, Δ risk-score vs. Δ marker) and -log p-value; colour denotes direction/significance (pink: significant negative; orange: non-significant negative; purple: non-significant positive; blue: significant positive). **G–L**. Translational applicability of MifRix in measuring therapy response via fecal microbiota transplantation (FMT). **G–I**. Δ GD MifRix-final-scores, responders vs. non-responders, across three FMT trials (DavarD_2023, KongL_2020, KumpP_2017); responders showed significantly greater GD-risk decreases (p≤0.05 for two cohorts, marginal p≤0.07 for KumpP_2017). **J.** In an independent Indian Crohn’s disease cohort (PRJEB11584), patients showed significantly higher GD risk than donors, reflecting disease activity. **K.** Greater improvement in Crohn’s Disease Activity Index (CDAI) correlated with significantly greater GD-risk decrease. **L.** Patients achieving remission showed significantly greater GD-risk decrease than non-remitters. Exact p-values are indicated for all pairwise boxplot comparisons.

We next probed whether MifRix could detect disease-associated microbial configurations at the precursor stage of disease, using colorectal polyps, a well-established CRC precursor lesion, as a test case. Applying MifRix to gut microbiomes from 117 individuals across three studies (**Table S14**), polyp-associated microbiomes consistently showed the highest predicted CRC risk (**Fig. 6B**), confirming premalignant lesions already harbour CRC-like microbial features. Additionally, these microbiomes predominantly clustered with CRC Signature 1 (**Fig. 6C**), the sub-signature linked to elevated T2D and CVD risk earlier in our investigation (**Fig. 5**), and showed correspondingly elevated CVD and T2D risk (**Fig. 6B**). These results reinforced CRC Signature 1 as an early microbial state in colorectal tumorigenesis, further confirming that MifRix’s cross-disease sub-signature associations reproduce robustly across independent datasets.

We next applied MifRix to the published NU-AGE Mediterranean-diet (MED-diet) study, which assessed dietary adherence in 612 non-frail/pre-frail older adults across five European countries, to test whether MifRix could measure intervention efficacy^77^. Across pre- and post-intervention timepoints, higher MED-diet adherence was negatively associated with risk for 7 of 10 diseases, significantly so for CRC, Schizophrenia, and MS (**Fig. 6D**), linking adherence to a healthier, disease-protective microbiome. MifRix risk-scores also tracked clinical markers: Parkinson’s risk decreases correlated with improved Constructional Praxis scores (**Fig. 6E**), while IBD risk correlated positively with IL-17, consistent with IL-17’s reported negative association with MED-diet-responsive taxa (**Fig. 6F**).

Finally, we tested whether MifRix risk-scores could monitor therapeutic response using three public Fecal Microbiota Transplantation (FMT) trials (**Table S14**). Across all investigated trials, patient gut microbiomes had the highest generic disease risk-scores at baseline (**Fig. S14**). Notably, responders showed significantly greater GD risk-score reductions than non-responders across all three cohorts (**Fig. 6G-I**), which was validated in an independent Crohn’s disease FMT cohort (PRJEB115184): healthy donors showed lower GD risk-scores than patients (**Fig. 6J**), post-FMT reductions correlated with CDAI improvement (**Fig. 6K**), and remitters showed greater risk decreases than non-remitters (**Fig. 6L**). Together, MifRix risk-scores identify disease-associated microbiomes and sensitively track clinical improvement during therapy.

## Discussion

The gut microbiome has emerged as a major contributor to human disease, yet most microbiome-based machine learning studies treat diseases as isolated, binary classification problems, overlooking the biological relationships among them. Here, we present MifRix, a unified framework that simultaneously estimates generic and disease-specific risk across ten major disorders by integrating taxonomic composition with microbiome-derived functional potential. Importantly, MifRix incorporates both sequencing strategy and industrialization status as model features, enabling platform-agnostic risk prediction across 16S rRNA and whole-genome shotgun datasets while accounting for population-level differences associated with industrialization. Beyond robust predictive performance across geographically and technologically diverse cohorts, MifRix links risk prediction to the specific taxa driving individual predictions and reveals substantial heterogeneity within disease-associated microbiome configurations, suggesting that microbiome-derived disease risk is best understood as a multidimensional continuum rather than a set of independent states.

A central finding is that microbiome-derived risk naturally recapitulates known inter-disease relationships, linking colorectal cancer with IBD and T2D, CVD with T2D, and multiple neurological disorders via shared neuroinflammatory pathways, despite no such relationships being explicitly encoded during training. Notably, 9 of the 10 disease-pairs exhibiting significant positive correlations in disease-specific risk-scores were also supported by previously reported disease co-occurrence, lending independent support to the inter-disease relationships captured by MifRix. The strong concordance between risk-score-based co-occurrence and microbial signature similarity supports the hypothesis that shared microbiome alterations contribute to the broader interconnected landscape of human disease.

A second advance is the demonstration that disease-associated microbiome alterations are themselves reproducibly heterogeneous. Using sample-level SHAP explanations, we show that each disease comprises multiple reproducible microbial sub-signatures, with cross-disease associations concentrated in specific sub-signatures rather than distributed uniformly across patients. This offers a plausible explanation for the limited cross-cohort transferability reported for prior microbiome classifiers, suggesting such heterogeneity is intrinsic to host-microbiome biology rather than purely technical or geographic in origin. This resolution also revealed that individual taxa, such as *Segatella copri*, can show disease-dependent, even opposing, associations across conditions, reinforcing that microbial contributions are context-dependent rather than fixed, and should be interpreted within disease-specific ecological networks rather than as universally ‘beneficial’ or ‘harmful’.

Translational analyses extend these findings beyond classification: elevated risk scores in clinically healthy individuals, CRC-like signatures in colorectal-polyp microbiomes, and risk-score reductions tracking clinical improvement after FMT and MED-diet intervention together suggest MifRix captures biologically meaningful, potentially pre-clinical microbiome remodelling. MifRix is trained on a largest, geographically and technologically diverse microbiome collections assembled to date, and includes dual taxonomic-functional integration, sample-level interpretability. Consistent performance across independent validation cohorts suggests that the framework captures robust biological signal rather than cohort-specific artifact.

However, here it is also important to highlight some of the limitations. The study remains retrospective, reliant on heterogeneous public datasets with variable clinical metadata, and, despite spanning 48 nationalities, still underrepresents non-industrialized populations; Prospective, longitudinal validation is also needed. Functional profiles were computationally inferred rather than directly measured, and disease categories (e.g., IBD, MDD) necessarily aggregate clinically heterogeneous subtypes, potentially masking finer-grained biological distinctions. Associations identified here are correlative, not causal, and roughly half of the significant disease-pairs identified lacked prior literature support, warranting independent replication. Finally, the current ten-disease scope, and the absence of host clinical and multi-omics data, both represent natural directions for expansion, alongside establishing an accessible, well-documented computational pipeline to support external validation and eventual clinical translation.

Despite this, MifRix extends microbiome-based disease prediction beyond binary classification, revealing shared and heterogeneous microbial architectures underlying human disease. As larger longitudinal microbiome datasets emerge, such frameworks may enable earlier risk assessment, refined patient stratification, and microbiome-informed therapeutic monitoring.

## Data availability

This paper analyzes existing, publicly available data, accessible at the European Nucleotide Archives (ENA) and curated Metagenomic Data (see **Methods**). Study details are listed in **Table S1**. However, the metadata for one of the cohorts used in this study (LifeLinesDeep_201695^78^) is subject to access restrictions and is not publicly available; this dataset can be accessed through the original publication (https://doi.org/10.1136/bmjopen-2014-006772). All the data generated in the form of results in our study have been deposited at https://github.com/Sourav-MiRe/Project_MifRix, and it is publicly available.

## Code availability

All original code has been deposited at https://github.com/Sourav-MiRe/Project_MifRix and the complete MifRix pipeline for generic and disease-specific risk prediction, identification of explainable features, and disease sub-signature prediction is publicly available at: https://github.com/Sourav-MiRe/MifRix.

## Acknowledgements

T.S.G. acknowledges the Department of Biotechnology, Ministry of Science and Technology, Government of India for the Ramalingaswami Re-entry Fellowship (BT/HRD/35/02/2006). T.S.G. acknowledges IIIT-Delhi for the Discovery Track Investigator grant. S.G. acknowledges IIIT-Delhi for the Institute-Fellowship.

## Author Contributions Statement

Conceptualization: T.S.G., S.G; Methodology, formal analysis, validation: S.G., T.S.G., N.A.; Investigation: S.G., T.S.G., N.A., P.S.; Data collection: S.G., A.A, D.P. (Debjit Pramanik), D.P. (Dinesh Palanimuthu), N.A.; Writing-original draft, T.S.G., S.G.; Writing-review and editing: T.S.G.; Supervision: T.S.G.; Funding acquisition: T.S.G.

## Methods

### Creating a global collection of high-quality genomes

We collected 1,52,063 genomes (including Metagenome Assembled Genomes or MAGs) encompassing 4,814 different microbial species-level taxa from different publicly available comprehensive databases like EMBL-EBI MGnify, UNINA-MAGs Genome Repository, NCBI-RefSeq^79–81^. Furthermore, to incorporate genomes from taxonomic lineages, typically lost with urbanization/industrialization, we also collected genomes from tribal communities of Nepal and Hadza hunter-gatherers of Tanzania (assembled and analyzed as part of previous study)^82^. To retain only the high-quality genomes, 71,238 genomes with completeness values >= 90 and contamination <= 5, were filtered out and considered for functional annotations.

### Genome annotation and creation of a species-level functional feature abundance profile

All 71,238 high-quality genomes were functionally annotated using eggNOG-mapper v2 against the eggNOG v5.0 database^83,84^. Briefly, genes were predicted using Prodigal^85^, translated into protein sequences, and annotated across eight functional categories: CAZy (carbohydrate-active enzymes), COG (Clusters of Orthologous Groups), BiGG, KEGG Orthologs, KEGG Reactions, KEGG Modules, EC (Enzyme Commission) numbers, and Pfam protein families. Genome-specific annotation files were merged using an in-house R script to generate genome-level functional abundance matrices based on the frequency of each functional feature.

The resulting matrix comprised functional abundance profiles for all 71,238 genomes (mean 15 genomes per species-level taxon). Genome-level profiles were subsequently aggregated to the species level by calculating the arithmetic mean abundance of each functional feature across all genomes within a species. The resulting taxa-to-function matrix contained mean abundances of 57,743 functional features across 4,814 species-level taxa. Inclusion of multiple genomes per species enabled measuring the prevalence of functional features across multiple strains of the same species-level taxa, capturing the confidence of detection of each functional feature across all genomes of the 4,814 species-level taxa. Biological descriptions for all functional feature identifiers were retrieved by web scraping the corresponding database interfaces using custom Python scripts, which were incorporated into our annotation pipeline. The resulting functional resource, termed the Functional Map, together with the associated scripts, has been deposited at https://github.com/Sourav-MiRe/Project_MifRix.

### Creating a global collection of gut microbiome profiles

Concomitantly, we created a global collection of 44,533 gut microbiomes from 189 study cohorts covering 48 countries and 6 continents around the globe encompassing 27,323 non-diseased individuals (referred to as ‘control’) and 17,210 microbiomes covering different diseases. The details of the 189 study cohorts and the distinct subsets of gut microbiomes (and study cohorts) included for model training, testing and validation (validation using unseen datasets and comparative validation with state-of-the-art indices) for different diseases as well as for validation in different intervention and population-level studies have been described in the **Table S1**. Of these, 18,751 (42%) were whole genome sequenced (WGS) microbiomes collected from 96 studies and 25,782 (58%) were 16S sequenced microbiomes collected from 93 studies.

### Data processing and generation of taxonomic abundance profiles

The species-level taxonomic abundance profiles of 62 of these study cohorts, already generated using metaphlan3^86^, (12,666 gut microbiomes) were directly available in the curatedMetagenomicData repository^87^ (see **Table S1**). For the rest of the WGS datasets, the same protocol as adopted by the curatedMetagenomicData was utilized using the updated version of metaphlan4^88^. All 16S derived gut microbiome datasets were downloaded from the corresponding data sources (as denoted by the accession number column in **Table S1**) processed using the SPINGO^89^ taxonomic classification tool. Metadata for datasets not included in curatedMetagenomicData were manually curated from the corresponding publications. To standardize taxonomic nomenclature across studies generated using different classification tools and database versions, all species-level taxa detected in the 44,533 microbiomes were then harmonized to the latest NCBI Taxonomy nomenclature using an in-house Python script. Taxa assigned multiple synonymous names were merged and renamed according to the updated NCBI taxonomy. The resulting initial abundance profile matrix (initial-AP) comprised 44,533 gut microbiomes from 189 study cohorts and 9,252 species-level taxa. As these taxa were detected in at least one microbiome, the matrix includes atypical or allochthonous species that may not represent common gut residents.

### Generation of microbiome-derived functional profile

The functional abundance profiles of 1,844 of these 9,252 species-level taxa were present in the list of 4,814 taxa, for which we generated the Functional Map as described above. Given the drastic reduction in taxa numbers, we then checked what fraction of the global microbiomes were represented by these 1,844 taxa. These 1,844 species-taxa accounted for a cumulative representation of at least 85% in 87% of the 44,533 gut microbiomes indicating that these capture the majority of the composition in our microbiome collection (**Fig. S1**).

Thus, in the next step, we reduced the number of taxa in our initial-AP to this list of 1,844 taxa and renormalized to generate our final taxonomic abundance profile containing the relative abundance of 1,844 taxa in 44,533 gut microbiomes. This was referred to as ‘AP’. Accordingly, we also reduced the Functional Map to these 1,844 taxa. Finally, we performed an inner matrix multiplication between the AP (44,533 gut microbiomes X 1,844 taxa) and the functional map (1,844 taxa X 57,743 functional features) to create a ‘Functional Profile’ matrix (referred to as ‘FP’) containing the abundances of 57,743 functional features, across the eight different functional categories, in 44,533 gut microbiomes (see **Fig. 1C**). This FP, along with the AP, served as a large-scale framework for leveraging microbiome-specific functions and abundances spanning different geographic locations, life-style and disease categories. Additionally, since sequencing type (16S or WGS) and cohort-population-type (Industrialized or Non-Industrialized) have been shown to have significant effects on the gut microbiome composition profiles in previous studies^90–93^, these two metadata columns were also added as digitized values in both the AP and FP.

### Model Development

#### Data pre-processing and Feature Selection

The next step was to build models leveraging the taxonomic abundance profiles (AP) as well as the microbiome-derived functional profiles (FP) for the prediction of generic disease-risk and the disease-specific risk-scores. For this purpose, we developed a ‘Two-Stage Risk-Score Prediction’ framework. The first step was to make a ‘Generic-Disease’ (GD) model to predict the probability of an individual having a generic disease-like microbiome signature capturing microbiome-alterations shared across multiple diseases. The next step was to devise models (disease-specific or DS) to predict disease-specific risk-scores. Different disease-specific models were developed for 10 different diseases, namely colorectal cancer (CRC), inflammatory bowel disorder (IBD), irritable bowel syndrome (IBS), cardiovascular disease (CVD), type-II diabetes (T2D), Parkinson’s disease (PD), multiple sclerosis (MS) and schizophrenia (SCZ), Alzheimer’s Disease (AD) and Major Depressive Disorder (MDD) where each model predicts the risk of one disease as compared to other nine diseases. These 10 different diseases had at least 300 gut microbiome profiles in our collection of 44,533.

Two variants of the GD model, MifRix-GD-AP and MifRix-GD-FP, were trained on 38,054 microbiomes from 150 cohorts (a subset of the 44,533), utilizing their taxonomic abundance profiles (AP) and microbiome functional profiles (FP) respectively. These included 24,114 microbiomes from control individuals (apparently non-diseased) and 13,940 microbiomes from individuals having at least one of the 10 different diseases we have considered. Of these microbiomes, 22,045 (58%) were 16S-based from 74 cohorts, while 16,009 (42%) were generated using WGS from 76 study-cohorts. For the GD model variants, all diseased microbiomes were labelled as class 1, while control samples were labelled as class 0.

Similarly, separate DS models were trained for each of the 10 diseases, with each DS model also having a taxonomy-based (AP) and microbiome-derived-function-based (FP) variants. Each DS model was trained excluding the controls, with the microbiomes of the target disease labelled as 1 and those of other nine diseases labelled as 0. The data utilized for training the DS models consisted of 13,940 microbiomes from diseased individuals encompassing 108 unique study-cohorts (**Table S2**). Among these microbiomes, 5,897 (42%) were generated using WGS from 39 cohorts, while 8,043 (58%) were 16S from 69 cohorts.

The approach for training each of the 22 models in the MifRix framework was the same as described below. The datasets were split into pure training (75% of the data) and model testing (25% of the data) sets using the ‘train_test_split’ function from python-based cuML package for efficient random sampling^94^. Features exhibiting near-zero variance in their abundance profiles across all training microbiome samples (variance threshold set at 10^-□^) were excluded from the analysis. To address the massive class imbalance in the datasets, a common challenge in biological datasets, random under-sampling was applied from the scikit-learn library on the training datasets to prevent bias toward the majority class during training of the models. Finally, to identify the most biologically relevant features for a model, a XGBoost-based feature selection was applied to each training dataset corresponding to the model to reduce dimensionality while preserving important features essential for accurate risk prediction. XGB run was performed 30 times and, in each iteration, features were ranked based on their importance scores using Gini impurity, and only those features with non-zero importance in at least 75% of the runs were retained for model training purposes. Moreover, sequencing strategy (16S rRNA or WGS) and industrialization status (industrialized or non-industrialized) of the microbiomes were included as additional features in each MifRix model to account for sequencing platform and population-level variation across cohorts.

### Development of the risk-score generation framework: Formulation of individuals models

To create an efficient, robust, and comprehensive risk-score generation framework, we leveraged 11 distinct machine learning models. Each model captures a unique perspective of the data, and their ensemble is expected to facilitate the identification of robust signatures by integrating these complementary viewpoints. The detailed approach has been mentioned in **Text S2**.

### Comparative analysis of the performance between MifRix-FP and MifRix-AP

MifRix-AP and MifRix-FP were compared both at the level of GD which is generic and also at the level of disease-specific DS models. Firstly, the GD models developed for both MifRix-AP and MifRix-FP were utilized to evaluate their performance in terms of AUC on test dataset (25% data; refer to **Fig. S2A-B**) and then the performance of GD-MifRix-AP and GD-MifRix-FP were compared on 32 completely unseen datasets. The evaluation was conducted based on accuracy to ensure a fair assessment of the models’ ability to distinguish between diseased and control microbiomes across diverse datasets (**Fig. S4A**). Also, the weighted mean accuracy was calculated for MifRix-AP, MifRix-FP and MifRix-final-score to show overall superiority towards generic disease-state forecasting. The next step was to evaluate the performance of disease-specific models (DS) using all three MifRix variants (MifRix-AP, MifRix-FP and MifRix-final-score) in the testing set (25% data) for each disease AUC was calculated using MifRix-FP and MifRix-AP and also the probability distributions of diseased microbiomes for each disease were compared using Wilcoxon signed-rank test in R (see **Figs. S2C-D**, **S3**). Finally, the performance of disease-specific MifRix-AP and DS-MifRix-FP were compared on 27 study-disease-pairs (25 study-cohorts out of which two studies namely LiJ_2014 (T2D, IBD), YuJ_2015 (T2D, CRC) have microbiomes from two disease conditions) using the study-wise recall, weighted mean recall across all the studies and the distribution of disease probabilities in each study (refer to **Fig. S4B**; **Table S3**).

### Comparative evaluation of the generic MifRix-GD ensemble predictors with the state-of-the-art microbiome health indices in terms of health status determination

The detailed approach of comparing the performance of the generic MifRix-AP and MifRix-FP based GD ensemble predictors with previously established microbiome-based health indices has been mentioned in **Text S3**.

### MifRix-based disease-specific risk assessment and capturing the inter-disease crosstalk

While the generic disease (GD) models detect microbiome alterations associated with disease, they primarily capture signatures shared across multiple conditions and therefore have limited disease specificity. To address this, we implemented disease-specific (DS) models that estimate susceptibility to each of the ten diseases by contrasting the target disease against the remaining nine using both taxonomic abundance and taxonomy-derived functional profiles.

DS models were applied to 3,675 microbiomes from the held-out testing set and 2,182 independent microbiomes from 25 validation cohorts encompassing all ten diseases. For each dataset, target disease risk scores were compared with the mean risk scores of the remaining nine diseases using boxplots to evaluate disease-specific discrimination.

Because related diseases may share microbiome alterations, we further investigated cross-disease associations using DS risk scores. For each target disease, microbiomes with risk scores >0.5 were selected. Within this subset, the number of microbiomes from each query disease with risk scores >0.5 was compared with those from all remaining diseases combined, generating pairwise disease enrichment tables. Association significance and direction were determined using Fisher’s exact test.

This analysis was performed independently for MifRix-AP, MifRix-FP, and the MifRix-final score across both testing and validation datasets. The MifRix-final score integrates AP and FP predictions to emphasize concordant signals while reducing the influence of discordant predictions. Disease associations were classified as positive or negative according to the Fisher’s exact test odds ratio.

Disease association networks were then constructed independently for the testing and validation datasets by integrating results from MifRix-AP, MifRix-FP, and the MifRix-final score. Only associations with consistent direction across all three scoring schemes were retained. Edge weights reflected reproducibility across the three frameworks and statistical significance (≤ 0.05): associations significant in all three frameworks were assigned weights of ±1, those significant in two frameworks ±0.7, and those significant in one framework ±0.3, with positive and negative values denoting enrichment and depletion, respectively.

Finally, only associations showing consistent directionality across both testing and validation networks were retained. These disease pairs were further filtered through literature curation, and only previously reported or biologically plausible associations were included in the final cross-disease microbiome interaction network (**Fig. 4G-H**).

### Adding explainability to the MifRix-derived Risk Scores using SHAP

MifRix incorporates an explainability module based on SHapley Additive exPlanations (SHAP) to quantify the contribution of individual features to each prediction, providing both global and local interpretability^95^. SHAP values were computed for the GD and DS MifRix-AP ensemble models using TreeExplainer or LinearExplainer, depending on the underlying algorithm. KernelExplainer was not applied to K-Nearest Neighbors and Naïve Bayes models because of its computational cost at this scale. Instead, SHAP values were generated for all supported ensemble components and aggregated across models using the median feature attribution.

### Shared microbial signatures and cross-disease similarity analyses

Disease risk scores generated by the MifRix-AP models were assembled into a unified disease-risk matrix. Pairwise disease associations were quantified using Spearman’s rank correlation, with significance calculated in R. To identify robust disease-associated microbial signatures, SHAP values from the disease-specific MifRix models were combined for each of the ten diseases. Within each disease, species were ranked by median SHAP value, and the top 50 positively contributing taxa were selected. Species with positive SHAP values in ≥70% of microbiomes within a disease were additionally retained, together with the union of top-ranked taxa across all diseases. For each disease, the relative abundances of the selected taxa were compared between target-disease microbiomes and all other diseases using two-sided Wilcoxon rank-sum tests, and effect sizes were estimated using Hedges’ g (R package effsize, v0.8.1). Taxa were classified as enriched or depleted according to the direction of the effect size and statistical significance (p ≤ 0.05, or p ≤ 0.10 to capture marginal associations), generating a disease-specific directionality matrix. Pairwise disease similarity was then quantified from this matrix using Manhattan distance and converted into a normalized microbial signature similarity matrix.

### Association between microbial signatures and disease co-occurrence

To determine whether shared microbial signatures reflected disease co-occurrence patterns, pairwise microbial signature similarities were compared with disease-disease associations derived from MifRix-AP risk-scores using Spearman’s rank correlation. The concordance between the two disease relationship matrices was further evaluated by principal coordinate analysis (PCoA) followed by a Procrustes randomization test with 999 permutations implemented in the ade4 package (v1.7-23). Scatter plots and ordination visualizations were generated using ggplot2 (v3.5.2) and ggrepel (v0.9.6).

### Disease-specific clustering of microbiomes based on DS risk-score-specific microbiome SHAP values and projection of independent validation microbiomes

To investigate whether disease-associated microbiomes comprise distinct microbial configurations, SHAP values from the disease-specific (DS) MifRix models were extracted for microbiomes predicted as disease-associated (risk score >0.5) in the held-out testing dataset. Independent validation microbiomes predicted as disease-associated and matching the target disease diagnosis were processed similarly and projected onto the disease-specific signatures derived from the testing cohort. For each disease, species with zero variance in the testing cohort were removed. SHAP values were standardized using the testing cohort mean and standard deviation, and the same scaling parameters were applied to the validation cohort. Principal component analysis (PCA) was performed on the standardized testing SHAP matrix, and validation microbiomes were projected into the testing-derived PCA space using the learned loading matrix without re-fitting the model.

To identify reproducible microbial signatures, the standardized testing SHAP matrix was subjected to k-means clustering, with the optimal number of clusters selected using the average silhouette coefficient. Final cluster assignments were obtained from a model fitted with 200 random initializations. Testing-derived signatures were visualized in PCA space, and multivariate normal confidence ellipses were constructed around each cluster. Validation microbiomes were projected into the same PCA space and assigned to disease-specific signatures based on inclusion within the corresponding confidence ellipses using a point-in-polygon algorithm. Microbiomes falling outside all ellipses were left unassigned. PCA, k-means clustering, silhouette analysis, and geometric projection were implemented using the R packages stats (v4.6.1), cluster (v2.1.8.2), sp (v2.2-1), ggplot2 (v4.0.3), dplyr (v1.2.1), and scales (v1.4.0).

### Identification of cluster-specific microbial signatures

To characterize the microbial composition of each disease-specific signature, the 20 species with the highest median positive SHAP values within each testing cluster were selected as candidate signature taxa. Testing and validation microbiomes assigned to the same cluster were independently compared with (i) non-diseased controls and (ii) microbiomes from all other diseases excluding the target disease. Differences in relative abundance were assessed using two-sided Wilcoxon rank-sum tests, and effect sizes were estimated using Hedges’ g. Taxa were classified as enriched or depleted according to effect size direction and statistical significance (Hedges’ g > 0 and p ≤ 0.05: +2; Hedges’ g > 0 and 0.05 < p ≤ 0.10: +1; Hedges ’g < 0 and 0.05 < p ≤ 0.10: –1; Hedges’ g < 0 and p ≤ 0.05: –2). These categorical scores were assembled into cluster-specific signature matrices and visualized as heatmaps to compare microbial enrichment and depletion patterns between the testing and validation cohorts. Statistical analyses were performed using the R packages effsize (v0.8.1), ComplexHeatmap (v2.18.1), circlize (v0.4.2), grid, and ggplot2.

### Association of microbial signatures with cross-disease risk profiles

To determine whether disease-specific microbial signatures were associated with distinct cross-disease susceptibility, microbiome-derived risk-scores from the testing and validation cohorts were merged according to their assigned disease signature. For each signature, MifRix-derived risk-scores corresponding to all ten diseases were compared across clusters using boxplots, allowing identification of microbial signatures exhibiting elevated risks toward additional diseases. Pairwise statistical comparisons between clusters were performed using two-sided Wilcoxon rank-sum tests after excluding clusters represented by fewer than five microbiomes. Visualization and statistical analyses were performed using ggplot2 (v4.0.3), dplyr (v1.2.1), reshape2 (v1.4.1), colorspace (v2.1.0) and RColorBrewer (v1.1.2).

### Clinical validation and translational evaluation of MifRix-derived disease risk-scores

To demonstrate the translational utility of MifRix, we evaluated the framework across nine independent cohorts (n = 1,905 microbiomes) encompassing diverse clinical applications. These analyses assessed disease risk prediction across ethnically distinct populations, detection of colorectal cancer-like signatures in premalignant colorectal polyps, associations between Mediterranean diet adherence and microbiome-derived disease risk, therapeutic response following fecal microbiota transplantation (FMT), and longitudinal changes in disease risk in relation to clinical outcomes in Crohn’s disease. Detailed analytical procedures and statistical methods are described in **Text S5**.

