## Supplementary Document for "MifRix: An Integrated microbiome framework for predicting generic and disease-specific risks investigating inter-disease diagnostic cross-talks and intra-disease signature variability"

### Text S1

#### Curation and functional annotation of microbial genomes for creating a species-level probabilistic functional profile

This Supplementary text documents the complete protocol utilized for the generation of species-level probabilistic functional profile. Since our objective was to also incorporate microbiome-level function profiles as a key component of our risk-prediction workflow, the subsequent requirement was to build a probabilistic association index of different functional annotations in each of the microbiome-associated taxa (identified in the microbiome taxonomic composition profiles described above). For this purpose, we meticulously curated 71,238 high-quality genomes, encompassing metagenome-assembled genomes (MAGs) as well as reference-/isolate-genomes from various sources, from 4,814 species-level microbial taxa, with an average genome count of 15 per taxon. These genomes were functionally annotated using an extensive combinatorial pipeline (see **Methods**) to generate, for each genome, an abundance-based feature vector of 57,743 functional "tags" spanning eight distinct functional categories: 129 carbohydrate-active enzymes (CAZy), 10,270 BiGG models (Biochemical, Genetic and Genomic knowledge-base), 11,588 clusters of orthologous groups (COGs), 3,255 enzyme classification numbers (EC), 10,641 protein families from InterPro, 734 KEGG Modules, 16,976 KEGG Orthologs, and 4,150 KEGG Reactions[1–6]. Collectively, this approach yielded a multi-layered functional characterization of the 71,238 microbial genomes spanning 4,814 species-level taxa (**Fig. 1A**; **Methods**).

This resulting matrix was then filtered to retain 1,844 species-level taxa specifically detected in the taxonomic composition profiles of the 44,581 gut microbiomes described above (hereafter referred to as Abundance-Profile, or AP). Collectively, these 1,844 taxa accounted for ≥85% relative abundance in 87% of our entire gut microbiome collection, and ≥70% relative abundance in 79% of the 150 study cohorts investigated here (**Fig. S1**). Among the 1,844 taxa identified, 33% belonged to the phylum Bacillota, 30% to Pseudomonadota, 15% to Actinomycetota, and 11% to Bacteroidota, making these the four most dominant phyla in terms of species representation (**Fig. 1B**). Of these 1,844 taxa, 53% had reference genomes, 23% had metagenome-assembled genomes (MAGs), and the remaining 23% had both reference genomes and MAGs. The genome-level multi-layered functional profiles were then aggregated to the species level by computing the mean prevalence of each functional feature across all genomes within a given species-level taxon, resulting in a species-resolved functional profile matrix.

### Text S2

### Design and Development of the MifRix Framework

There were 22 models trained and developed within the MifRix workflow: One set of models for the taxonomic abundance based MifRix-AP variant and another set of models based on taxonomic-composition-informed functional abundance variant MifRix-FP. Each set further comprising 11 models: one GD model for generic-disease-risk, 10 disease-specific (DS) models for measuring risks for the 10 diseases.

A key question in the design of the MifRix framework was the choice of the machine-learning approach for predicting risk-scores. A key consideration in this selection is the inherently complex and interactive nature of microbial communities. The gut microbiome is not a simple taxa collection but a dynamic ecosystem shaped by interactions among microbes and between microbes and the host environment, spanning both taxonomic and functional layers. These involve intricate, non-linear relationships that are unlikely to be captured by a single machine-learning approach. To address this complexity, we adopted an ensemble machine-learning strategy.

Each of the 22 models was trained, following hyperparameter optimization, using an ensemble of 11 different machine-learning algorithms integrated through a voting-based predictor. The optimal method/algorithm combination was then selected to maximize predictive performance and robustness (**Fig. 2A**; Refer to **Methods**). This approach is described below:

#### ***Detailed description of the training and development of the ensemble machine learning approach of MifRix***

To create an efficient, robust, and comprehensive risk-score generation framework, we leveraged 11 distinct machine learning models. Each model captures a unique perspective of the data, and their ensemble is expected to facilitate the identification of robust signatures by integrating these complementary viewpoints. This approach was further strengthened by using a large training dataset.

Three interpretable baseline approaches, namely Logistic Regression, Naïve Bayes, and K-Nearest Neighbors (KNN), were included for their ability to provide transparent risk estimates and capture foundational patterns in sparse microbiome data. Logistic Regression estimates disease probability, Naïve Bayes operates under feature-independence assumptions, and KNN infers disease risk based on similarity to neighbouring samples. Additionally, since taxonomic and functional abundance profiles (represented by AP and FP, respectively) are reflections of complex, non-linear relationships between taxa and/or their functions, ensemble models are required to accurately capture these intricacies. To capture the complex, non-linear relationships within microbiome data, we incorporated eight tree-based ensemble approaches, including Random Forest, Extremely Randomized Trees (Extra Trees), Gradient Boosting, HistGradientBoosting, XGBoost, LightGBM, Bagging Classifier, and CatBoost[7–15]. These employ bagging or boosting strategies to improve predictive performance, reduce overfitting, and iteratively refine decision boundaries, making them well-suited for high-dimensional microbiome datasets. Each approach thus captures a unique perspective of the data, and their ensemble is expected to facilitate the identification of robust signatures by integrating these complementary viewpoints.

The combinatorial workflow utilized a pre-training phase of feature selection, random undersampling and hyperparameter tuning to ensure efficient and reliable performance. The hyper-parameter tuning was using Optuna[16]. Optuna is a Bayesian optimization framework that efficiently explores the hyperparameter space to find optimal hyperparameters. Candidate models were hyperparameter-tuned (with the objective of prioritizing recall within the recall-specificity trade-off.) using a composite objective function ($0.7 \times$ recall $+ 0.3 \times$ specificity), and those achieving specificity >= 60% and AUC > 90% were selected for inclusion in the final ensemble predictor. Given the significance of accurately assessing disease risk, recall was emphasized to capture as many at-risk individuals as possible, minimizing the risk of false negatives while keeping false positives at a reasonable level. Considering the huge variation and sparsity in the microbiome data, this recall-specificity trade-off is highly needed.

Finally, give the complex and non-linear relationships within microbiome data, we hypothesized that no single model could fully capture the patterns of microbiome variability. Therefore, we developed a voting-based risk-scoring system by combining multiple models. This system leveraged the median of probability outputs from different combinations of the 11 different ML models (as described below) and subsequently found the best combination (using the approach described below). This combination was then utilized to generate the risk-score. This approach is expected to effectively recognize microbiome variability patterns and ensure the generation of robust and reliable risk-scores. Since there were 11 different ML models, all combinations of model ensembles ($2^{11}$ - 1) were evaluated by taking the median (to reduce the impact of outliers) of their probability outputs. Subsequently, ensembles that achieved AUC > 90% and specificity

>= 60% were first selected and amongst them, the one with the highest recall was selected as the best combination (**Fig. 2B**).

The model-building process, for the GD and DS models, was performed using a subset of 38,054 and 13,940 microbiomes (75% training and 25% testing) selected from our global 44,533 microbiome collection (see **Methods** for details; **Table S2** for the model-building training and testing subsets utilized for the GD model and each of the 10 DS models).

Additionally, we incorporated an Explainability Module into the MifRix workflow, built upon Shapley Additive Explanations (SHAP). This module quantifies the contribution of individual features to the prediction generated by each model for a given microbiome sample and is described in detail in a subsequent section. For each of the 11 models (the Generic-Disease model and 10 Disease-Specific models), we additionally computed a MifRix-final-score by integrating the risk scores from the MifRix-AP and MifRix-FP variants. The MifRix-final-score was calculated as the mean of the MifRix-AP- and MifRix-FP-derived risk scores multiplied by (1 − Gini), thereby down-weighting microbiomes exhibiting greater inequality between the two risk-score variants (**Fig. 2C**).

### Text S3

**Comparative evaluation of the generic MifRix-GD ensemble predictors with the state-of-the-art microbiome health indices in terms of health status determination**

To compare the performance of the generic MifRix-AP and MifRix-FP based GD ensemble predictors with previously established microbiome-based health indices, namely the Gut Microbiome Health Index (GMHI), Gut Microbiome Wellness Index (GMWI 2), HACK-Top17-Score (the score associated to top 17 taxa from the ranking order of 201 Health-Associated Core Keystone taxa) and the Dysbiosis Score, we utilized eight WGS and twenty 16S 'unseen' study-cohorts[17–20]. Each of the datasets included microbiomes from both control and diseased individuals and had not been used during the training of any of the models in MifRix. GMHI is a microbiome-based binary classification score that uses species-level relative abundances to estimate host wellness, while GMWI 2 extends this by incorporating updated taxonomic curation and revised healthy vs. disease species references and using a logistic regression model but both GMHI and GMWI 2 have the limitation that only WGS datasets can be processed through them. HACK-Top17-score is generated as the summation of rank scaled HACK scores of the top 17 taxa which are present in a microbiome to indicate how healthy a particular microbiome is. The Dysbiosis Score, in contrast, measures microbial imbalance by using a Bray-Curtis distance-based approach to compare the microbial composition of individual samples against a healthy baseline, in our case the matched controls.

To do a thorough comparison with these methods, we utilized two approaches. First, we used a statistical approach based on the Mann-Whitney U test to determine the significant associations of each scores' distribution between control and disease groups. For each dataset, we computed index-wise p-values using the Wilcoxon rank-sum test, along with the direction of the mean shifts and based on the associations and significance three categories of associations (the mean probability across the diseased samples is significantly (taking a soft threshold of p-value $<= 0.1$ through Mann-Whitney test) higher than the mean probability across the control samples, the mean probability across the control samples is significantly (taking a soft threshold of p-value $<= 0.1$ through Mann-Whitney test) higher than the mean probability across the diseased samples and the final category is where there is no statistical significance of association (p-value $> 0.1$). As MifRix-AP, MifRix-FP, MifRix-final-score and Dysbiosis Score generate the scores to predict the disease status, ideally these three indices should be inclined towards higher mean probability across the diseased samples and on the other hand, GMHI, GMWI 2 and HACK-Top17-Score indicates the score towards wellness (measurement of healthy gut microbiome) so these three indices should be ideally inclined towards higher mean probability across the control samples. The percentage of studies where each method is showing correct significant association was calculated to compare the overall discriminatory power (see **Fig. 3A**). Second, we employed SHAP (SHapley Additive exPlanations) analysis within a LGBM classification model to determine how much each score contributed to model predictions when considered alongside other scores. We chose LGBM due to its effectiveness in handling data with high accuracy, its robustness to feature correlation, and its compatibility with tree-based SHAP explanation. Specifically, for each cohort, we trained an LGBM classifier using all scores simultaneously and computed the median absolute SHAP value per method. Finally, the six methods were ranked based on the median SHAP values for each study-cohort, which allowed us to evaluate the additive contribution of each method's score toward final prediction in a model-agnostic, interpretable way (see **Fig. 3B**). By leveraging these three approaches where we focus on feature importance on a tree-based classifier, statistical separability between distributions and model-aware feature attribution, we evaluate and compare the discriminative strength and statistical robustness of our MifRix ensemble predictors against established microbiome health indices across diverse unseen datasets.

### Text S4

#### Each Disease Is Characterized by Multiple, Distinct Microbiome Sub-Signatures

A single disease can arise through multiple microbiome configurations, each represented by a distinct microbial signature despite sharing the same clinical phenotype. Using SHAP values derived from the disease-specific MifRix models, we identified multiple disease sub-signatures within each disorder that were characterized by unique taxonomic directionality (enrichment and depletion) patterns (see **Figs. S10-S13**; **Fig. 5**; **Methods**). These signature-specific microbial alterations were highly reproducible when compared with both healthy controls and microbiomes from other diseases and remained stable following projection of completely independent validation cohorts onto the signature space derived from the testing dataset. Importantly, individual microbial signatures were associated with distinct cross-disease risk profiles, indicating that disease co-occurrence is driven by specific microbial configurations rather than by a single canonical disease microbiome. The current text details these sub-signatures for each of the 10 diseases as well as the taxa alterations that are characteristic of each sub-signature.

We began with IBD, for which four reproducible microbial signatures were identified, with testing and validation microbiomes each mapping onto the four signatures at varying frequencies. IBD-signature-1 was characterized by enrichment of *Collinsella aerofaciens*, and depletion of *Mediterraneibacter glycyrrhizinilyticus* and *Blautia schinkii*. IBD-signature-2 was primarily characterized by depletion of several short-chain fatty acid (SCFA)-producing commensals, including the butyrate-associated taxa *Gemmiger formicilis*, *Agathobaculum desmolans*, *Mediterraneibacter glycyrrhizinilyticus* and *Blautia* (B. glucerasea/schinkii), together with depletion of *Oscillibacter valericigenes* and the propionate producer *Phascolarctobacterium faecium*, whereas IBD-signature-3 showed enrichment of *Anaerostipes hadrus* accompanied by depletion of *Allocoprococcus comes*, *Fusicatenibacter saccharivorans*, *Blautia wexlerae* and *Streptococcus salivarius*. In contrast, IBD-signature-4 was distinguished by depletion of *Ruminococcus bromii*, *Anaerobutyricum hallii* and *Bifidobacterium pseudocatenulatum*. Despite of having a mixture of 16S and WGS microbiomes, these microbial configurations were consistently reproduced across both cohorts. Notably, individuals assigned to both IBD-signature-1 and IBD-signature-2 exhibited elevated predicted risks of both CRC and T2D. This increased cross-disease risk was accompanied by shared depletion of taxa including *Oscillibacter valericigenes* and *Blautia obeum* across IBD-signature-2, CRC and T2D, together with depletion of *Acetivibrio thermocellus* shared with CRC and depletion of *Phascolarctobacterium faecium* and *Blautia glucerasea* shared with T2D, suggesting that these common microbial alterations may underlie the observed disease associations[21–24] (**Fig. S10**).

Three distinct microbial alteration signatures were identified for CRC, with the vast majority (96.1%) of independent validation microbiomes, encompassing both 16S and WGS sequencing type, mapping to CRC-signature-1, indicating that this represents the predominant microbial configuration underlying CRC. CRC-signature-1 was characterized by enrichment of *Ruminococcus torques* and *Bacteroides uniformis*, together with depletion of several health-associated commensals, including *Oscillibacter valericigenes*, *Sutterella wadsworthensis*, *Subdoligranulum variabile*, *Blautia* (B. obeum/faecis) and *Dialister invisus*. In contrast, CRC-signature-2 exhibited enrichment of opportunistic oral- and inflammation-associated taxa, including *Porphyromonas asaccharolytica*, *Fusobacterium periodonticum* and *Phascolarctobacterium faecium*, whereas CRC-signature-3 was distinguished by enrichment of well-established CRC-associated pathobionts, including *Bacteroides fragilis*, *Parvimonas micra* and *Peptostreptococcus stomatis*. Rather than representing a single canonical CRC microbiome, these well-established CRC-associated taxa define multiple, distinct microbial sub-signatures, underscoring the existence of heterogeneous microbial configurations associated with the disease[25–28]. Notably, microbiomes assigned to CRC-signature-1 exhibited significantly elevated predicted T2D and CVD risk, suggesting that this microbial subtype may represent a shared microbial state linking CRC and metabolic dysfunction. This association was accompanied by common depletion of *Oscillibacter valericigenes*, *Dialister invisus* and *Blautia* (*B. obeum*/*faecis*) across CRC-signature-1 and T2D, implicating these shared microbial alterations as potential drivers of the observed disease co-occurrence (**Figs. S10B-C, S11A**). CRC-signature-3 was additionally associated with higher CVD risk.

Three reproducible microbial alteration signatures were also identified for CVD, with all independent validation microbiomes mapping to CVD-signature-1, indicating that this represents the predominant microbial configuration associated with cardiovascular disease. CVD-signature-1 was characterized by enrichment of several inflammation-associated taxa, including *Megamonas funiformis*, *Phocaeicola* (*P. plebeius*/*coprocola*), *Segatella copri* and *Acetivibrio thermocellus*, together with depletion of *Allocoprococcus comes* and *Schaalia odontolytica*. In contrast, CVD-signature-2 was distinguished by enrichment of opportunistic taxa, including *Escherichia coli*,

*Klebsiella pneumoniae*, *Gemella sanguinis*, *Veillonella parvula*, *Agathobaculum butyriciproducens* and *Ruthenibacterium lactatiformans*, including the oral-associated species *Gemella sanguinis* and *Veillonella parvula*. CVD-signature-3, on the other hand, exhibited enrichment of *Lawsonibacter asaccharolyticus* and *Turicimonas muris* together with depletion of beneficial commensals, including the probiotic species *Bifidobacterium longum* and the lactic acid bacterium *Lactococcus lactis*. Individuals assigned to CVD-signature-1 exhibited markedly elevated predicted risks of both IBS and SCZ, supporting a shared gut-cardio-brain microbial axis. This relationship was accompanied by common enrichment of *Phocaeicola* (P. plebeius/coprocola) across CVD, IBS and SCZ, together with shared enrichment of *Megamonas funiformis* and *Segatella copri* and depletion of *Schaalia odontolytica* between CVD and schizophrenia, suggesting that these conserved microbial alterations may contribute to the observed co-occurrence of cardiovascular, gastrointestinal and neuropsychiatric disorders[29–37] (**Fig. S11**). On the other hand, CVD-signature-1 and -2 showed higher risk of gastrointestinal-linked/metabolic disorders, CRC and T2D (CVD-signature-2 to both CRC and T2D and CVD-signature-3 to T2D) (refer to **Figs. 5, S11A, S10B-C**).

Microbiome alterations in T2D segregated into three reproducible microbial signatures, with 99.5% of independent validation microbiomes mapping to T2D-signature-1, indicating that this represents the predominant microbial configuration associated with type-II diabetes. T2D-signature-1 was characterized by enrichment of facultative and inflammation-associated taxa, including *Streptococcus salivarius*, *Escherichia coli*, and *Ruthenibacterium lactatiformans*, accompanied by depletion of several health-associated commensals, including *Oscillibacter valericigenes*, *Blautia* (B. glucerasea/obeum/faecis), *Dialister invisus* and *Phocaeicola coprocola*, reflecting a profound disruption of the beneficial gut microbial community. In contrast, T2D-signature-2 was distinguished by enrichment of the opportunistic bacteria *Aerococcus viridans* and *Enterococcus hirae*, whereas T2D-signature-3 exhibited enrichment of fermentative taxa, including *Gemmiger formicilis*, together with depletion of beneficial *Bifidobacterium* species, suggesting an alternative dysbiotic microbial configuration. Approximately half of the individuals assigned to T2D-signature-1 exhibited predicted CRC risk above the classification threshold, while a substantial proportion also displayed elevated CVD risk, indicating that this microbial subtype represents a shared microbial state linking metabolic, inflammatory and cardiovascular diseases. These cross-disease associations were accompanied by common depletion of *Blautia obeum* across

T2D, CRC and CVD, shared depletion of *Oscillibacter valericigenes*, *Dialister invisus* and *Blautia* (B. faecis/obeum) between T2D and CRC, and shared enrichment of *Ruthenibacterium lactatiformans* and *Escherichia coli* between T2D and CVD, suggesting that these conserved microbial alterations may contribute to the observed overlap between metabolic, inflammatory and cardiovascular disease states[38–48] (**Figs. S10B-C, S11A**).

The observed association between CVD microbial signatures and elevated risks of IBS and schizophrenia suggested that shared microbial configurations may extend beyond inflammatory and cardio-metabolic diseases into disorders of the gut-brain axis. We therefore investigated disease-specific microbial signatures across IBS, Alzheimer's disease (AD), schizophrenia (SCZ), Parkinson's disease (PD), multiple sclerosis (MS) and major depressive disorder (MDD), revealing reproducible microbial configurations that explain disease heterogeneity and shared cross-disease associations.

Among the three microbial signatures identified for IBS, IBS-signature-2 and IBS-signature-3 accounted for the entire independent validation cohort, comprising 40.7% and 59.3% of validation microbiomes, respectively, suggesting that these represent the predominant microbial configurations associated with IBS. IBS-signature-1 was characterized by enrichment of *Pseudocoprococcus catus*, *Alistipes* (*A. communis/onderdonkii*) and *Streptococcus salivarius*, together with depletion of several beneficial commensals, including *Roseburia faecis*, *Gemmiger formicilis*, *Flavonifractor plautii*, *Fusicatenibacter saccharivorans* and *Intestinimonas butyriciproducens*, indicative of reduced short-chain fatty acid-producing capacity. In contrast, IBS-signature-2 exhibited enrichment of inflammation-associated taxa, including *Escherichia coli*, *Phocaeicola* (*P. plebeius/coprocola*) and *Clostridium polysaccharolyticum*, accompanied by depletion of *Butyrivibrio fibrisolvens*, *Agathobaculum butyriciproducens* and other health-associated commensals, whereas IBS-signature-3 was primarily distinguished by depletion of *Papillibacter cinnamivorans*, *Collinsella aerofaciens*, *Lachnospira pectinoschiza* and *Ruthenibacterium lactatiformans*. Notably, microbiomes assigned to IBS-signature-2 exhibited progressively elevated predicted risks of CVD, AD and SCZ, suggesting that this microbial subtype represents a shared gut microbial configuration linking gastrointestinal, cardiovascular and neurological disorders. This was also reflected in terms of shared microbiome signatures, where in IBS-signature-2 shared enrichment of *Phocaeicola* (*P. plebeius/coprocola*) and *Escherichia coli* with CVD, enrichment of *Escherichia coli* and *Clostridium polysaccharolyticum* with AD, and enrichment of *Escherichia coli* together with *Phocaeicola* (*P. plebeius/coprocola*) with SCZ, providing a potential microbial basis for these cross-disease associations[29,49–56] (**Figs. S11, S12A**).

Alzheimer's disease (AD) exhibited four distinct microbial alteration signatures; however, AD-signature-1 represented the predominant microbial configuration, encompassing 95.7% of the independent validation microbiomes. This signature was characterized by enrichment of several opportunistic and inflammation-associated taxa, including *Escherichia coli*, *Klebsiella pneumoniae*, *Schaalia odontolytica*, *Clostridium scindens*, *Anaerotruncus colihominis*, *Bifidobacterium* (*B. dentium/pseudocatenulatum*), *Phascolarctobacterium faecium* and *Lancefieldella parvula*, together with depletion of health-associated commensals such as *Lachnospira pectinoschiza*, *Amedibacillus dolichus*, *Holdemanella biformis* and *Enterocloster clostridioformis*. In contrast, AD-signature-2 was enriched in metabolically active commensals, including *Eubacterium coprostanoligenes*, *Christensenella minuta*, *Dialister succinatiphilus*, *Oxalobacter formigenes* and *Clostridium polysaccharolyticum*, whereas AD-signature-3 displayed enrichment of *Ruthenibacterium lactatiformans*, *Streptococcus* (*S. salivarius/anginosus*) together with *Mogibacterium diversum* and *Evtepia gabavorous*. AD-signature-4, on the other hand, was distinguished by enrichment of *Faecalicatena contorta*, *Amedibacillus dolichus*, *Porphyromonas catoniae* and *Lactonifactor longoviformis*, representing an alternative microbial configuration within AD. Notably, individuals assigned to AD-Signature1 exhibited elevated predicted CVD and IBS risk, suggesting a shared microbial state linking neurodegeneration and cardiovascular disease. This relationship was accompanied by common enrichment of *Escherichia coli*, *Klebsiella pneumoniae* and *Phascolarctobacterium faecium* between AD-signature-1 and other two diseases, supporting the notion that expansion of opportunistic pathobionts and pro-inflammatory microbial taxa may contribute to the well-recognized association between cardiovascular dysfunction and Alzheimer's disease[50,56–61] (**Figs. S12A, S11A-B**). AD-signature-2 also showed relatively higher risk-scores for CVD. In contrast, AD-signature-3 and AD-signature-4 showed elevated risk-scores for MDD and MS (**Figs. S12A-B**, **S13A**)

Schizophrenia (SCZ) displayed four reproducible microbial alteration signatures. SCZ-signature-1 exhibited enrichment of *Escherichia coli*, *Bilophila wadsworthia* and *Bacteroides stercoris*, accompanied by depletion of the health-associated commensals *Adlercreutzia equolifaciens* and *Fusicatenibacter saccharivorans*. SCZ-signature-2 and SCZ-signature-4 accounting for the majority (62.8% and 26.4%, respectively) of the independent validation microbiomes, indicating that these represent the predominant microbial configurations associated with schizophrenia. Consistent with the shared gut-cardio-brain axis identified above, microbiomes assigned to SCZ-signature-2 (which contains both 16S and WGS microbiomes) exhibited markedly elevated predicted CVD risk and recapitulated the microbial alterations previously observed in CVD-signature-1. Specifically, SCZ-signature-2 was characterized by enrichment of inflammation-associated taxa, including *Megamonas funiformis*, *Phocaeicola* (*P. plebeius/coprocola*), *Leyella stercorea*, *Segatella copri*, *Bilophila wadsworthia* and *Fusobacterium mortiferum*, together with depletion of *Haemophilus parainfluenzae* and *Fusicatenibacter saccharivorans*, suggesting a pro-inflammatory microbial configuration[62–71]. Elevated risk for CVD was also observed for SCZ-signature-3. SCZ-signature-3 was distinguished by enrichment of *Megasphaera elsdenii*, *Lancefieldella* (*L. parvula/rimae*), *Pelagibacterium halotolerans* and *Schaalia odontolytica*. SCZ-signature-4, on the other hand, was characterized by depletion of several beneficial taxa, including *Schaalia odontolytica*, *Clostridium leptum*, *Dorea longicatena*, *Streptococcus thermophilus* and *Faecalibacterium prausnitzii*[50,57,67,68,70–74]. Beyond the CVD association, elevated predicted PD risk was consistently observed across SCZ-signatures-1 and -2. This relationship was accompanied by shared enrichment of *Bilophila wadsworthia* and *Escherichia coli*, together with depletion of *Faecalibacterium prausnitzii*, *Adlercreutzia equolifaciens* and *Haemophilus parainfluenzae*, suggesting that disruption of anti-inflammatory commensals alongside expansion of pro-inflammatory pathobionts may contribute to the microbial overlap between schizophrenia and Parkinson's disease[68,70,75,75–78] (**Figs S11A, S11C, S13B**).

Parkinson's disease (PD) was resolved into three reproducible microbial signatures, with PD-signature-3 representing the predominant microbial configuration, encompassing 99.1% of the independent validation microbiomes. This signature was characterized by enrichment of taxa implicated in host immune modulation and mucosal metabolism, including *Streptococcus thermophilus*, *Bilophila wadsworthia*, *Akkermansia muciniphila*, *Blautia schinkii*, *Flavonifractor plautii*, *Oscillibacter valericigenes*, *Sporobacter termitidis* and *Christensenella minuta*. In contrast, PD-Signature1 exhibited enrichment of *Alistipes communis* and *Lachnospira pectinoschiza*, together with depletion of *Blautia luti*, *Thomasclavelia ramosa*, *Turicimonas muris*, *Anaerobutyricum hallii*, *Dorea formicigenerans* and *Gordonibacter pamelaeae*, suggesting reduced abundance of several health-associated commensals. PD-signature-2, on the other hand, was distinguished by enrichment of *Amedibacillus dolichus*, *Anaerostipes hadrus*, *Blautia* (*B. obeum/faecis*) and lactic acid bacteria including *Latilactobacillus sakei* and *Leuconostoc mesenteroides*, accompanied by depletion of *Mediterraneibacter gnavus*, *Segatella copri*, *Faecalibacterium prausnitzii*, *Bacteroides* (*B. uniformis/xylanisolvens*), *Phocaeicola vulgatus*, *Bifidobacterium dentium* and *Adlercreutzia equolifaciens*. Notably, individuals assigned to PD-signature-3 exhibited markedly elevated predicted MS risk, consistent with the reciprocal disease association observed for MS. This overlap was accompanied by shared enrichment of *Streptococcus thermophilus* and *Blautia schinkii* between PD and MS, suggesting that these conserved microbial alterations may contribute to the well-recognized immunological and neuroinflammatory links between the two neurodegenerative disorders[79,80] (**Fig. S13**).

Unlike the other diseases investigated, MS exhibited substantial microbial heterogeneity, with three major microbial signatures collectively accounting for nearly all testing and validation microbiomes. MS-signature-2, MS-signature-3 and MS-signature-4 comprised 49.8%, 17.6% and 30.4% of the testing microbiomes, respectively, and similarly represented 28.0%, 18.5% and 53.6% of the independent validation cohort, demonstrating that these heterogeneous microbial configurations were consistently reproduced across datasets. MS-signatures-2 and -4 shared a common microbial architecture characterized by enrichment of *Mediterraneibacter glycyrrhizinilyticus*, *Streptococcus thermophilus*, *Agathobaculum desmolans* and *Blautia* (*B. obeum/glucerasea/schinkii/faecis*), suggesting a conserved microbial configuration underlying a substantial proportion of MS microbiomes. In contrast, MS-signature-2 was further distinguished by enrichment of *Faecalispora sporosphaeroides* and *Blautia* (*B. wexlerae/stercoris*), together with depletion of *Segatella copri*, *Bacteroides clarus* and *Leyella stercorea*, whereas MS-signature-3 exhibited a markedly different microbial composition, representing an alternative disease-associated microbial state. Consistent with the reciprocal relationship identified for PD, individuals assigned to MS-signature-3 and MS-signature-4 exhibited elevated predicted PD risk. This association was accompanied by shared enrichment of *Streptococcus thermophilus* and *Blautia schinkii*, reinforcing the existence of a conserved microbial signature linking these two neuroinflammatory disorders and providing further support for the microbiome-mediated overlap between PD and MS[79,80] (**Fig. S13**).

Finally, MDD was characterized by three microbiome signatures, with MDD-signature-1 alone accounting for 27.6% of testing microbiomes and 89.3% of the independent validation cohort, indicating that this signature captures the dominant microbial landscape associated with depression. MDD-Signature1 was characterized by enrichment of opportunistic and inflammation-associated taxa, including *Escherichia coli*, *Mogibacterium diversum*, *Streptococcus* (*S. salivarius/anginosus*), *Bifidobacterium* (*B. longum/pseudocatenulatum*), *Dorea formicigenerans* and *Blautia obeum*, together with depletion of several health-associated commensals, including *Faecalibacterium prausnitzii*, *Dialister invisus*, *Intestinimonas butyriciproducens*, *Enterocloster clostridioformis*, *Holdemanella biformis* and *Blautia faecis*, reflecting a microbial configuration consistent with intestinal dysbiosis and loss of anti-inflammatory taxa. Individuals assigned to MDD-Signature1 exhibited elevated predicted AD risk, supporting the close microbial relationship between depression and neurodegeneration. This association was accompanied by shared enrichment of *Escherichia coli*, *Mogibacterium diversum*, *Streptococcus* (*S. salivarius/anginosus*) and *Bifidobacterium pseudocatenulatum*, together with shared depletion of *Holdemanella biformis*, suggesting that these conserved microbial alterations may contribute to the well-established association between major depressive disorder and Alzheimer's disease through the gut-brain axis[54,81–84] (**Fig. S12**). MDD-signature-2 was associated with increase of *Bifidobacterium pseudocatenulatum, Alistipes communis, Evtepia gabavorous* and decrease of *Streptococcus salivarius* and *Blautia faecis*. MDD-signature-2 was also associated with increased risk of AD and SCZ. This association was accompanied by the common enrichment of *Bifidobacterium pseudocatenulatum* in both AD and SCZ and a common enrichment of *Evtepia gabavorous* in AD (**Fig. S11C, S12**). Moreover, MDD-signature-3 was, on the other hand, associated with an enrichment of *Nitrospira japonica, Clostridium clostridioforme 2 1 49FAA, Eubacterium sulci, Roseburia inulinivorans, Parasutterella excrementihominis* and depletion of *Helicobacter pylori, Lawsonibacter asaccharolyticus* and *Streptococcus anginosus*. This signature was associated with elevated risk of AD and MS (**Figs. S12, S13A**).

Collectively, these findings demonstrate that 10 diseases could be arranged as spectrum based on their mutual association patterns, with gastro-intestinal linked, metabolic disorders, gradually progressing neurological disorders at the other end of the spectrum. However, this granular investigation reveals that each disease comprises by multiple reproducible microbiome alteration signatures, with specific signatures mediating shared microbial alterations that mirror cross-disease risk relationships and provide a potential microbial basis for the interconnected nature of gastrointestinal, neurological and neuropsychiatric disorders.

### Text S5

**Clinical validation and translational evaluation of MifRix-derived disease risk-scores**

To capture the translational potential of MifRix-derived risk-scores we validated diverse utilities of the framework using 1905 microbiomes across nine independent cohorts (**Tables S1, S14**). To assess the robustness of disease-specific risk prediction across ethnically diverse populations, the MifRix-final-score for Type-II Diabetes (T2D) was evaluated using the independent Dwiyanto_2023 cohort comprising Chinese, Indian, Jakun and Malay participants[85]. Since information on obesity and elevated blood pressure was available for this cohort, predicted T2D risk-scores were compared across ethnic groups to examine whether MifRix captured differences in metabolic disease susceptibility. Pairwise group comparisons were performed using Dunn's multiple comparison test following Kruskal-Wallis analysis to check the statistical significance.

Next, to investigate whether premalignant colorectal polyps exhibit microbial configurations resembling colorectal cancer, MifRix-final-scores for CRC were generated for microbiomes from individuals with colorectal polyps obtained from three independent studies (FengQ_2015, YachidaS_2019 and ZellerG_2014). The distribution of predicted disease risk across all MifRix disease models was examined. To determine whether polyp microbiomes resembled specific CRC microbial configurations, microbiomes were projected onto the principal component space generated from the CRC microbial sub-signatures identified by MifRix. Their proximity to individual CRC sub-signatures was subsequently evaluated relative to both the testing dataset and completely independent CRC validation microbiomes.

For evaluating the clinical utility of MifRix-derived risk-scores in an independent population with diet intervention, we assessed the effect of MED-diet adherence on the gut microbiome of 612 non-frail/pre-frail older adults across five European countries[86]. Disease risk scores generated by the MifRix framework, encompassing Alzheimer's disease (AD), colorectal cancer (CRC), cardiovascular disease (CVD), inflammatory bowel disease and other gut inflammations (IBD+), irritable bowel syndrome (IBS), multiple sclerosis (MS), Parkinson's disease (PD), schizophrenia (SCZ), type 2 diabetes (T2D), and major depressive disorder (MDD) and generic disease risk (GD) were used to compute a cumulative disease risk score for each participant, defined as the number of diseases where risk score > 0.5 for an individual. To assess the relationship between Mediterranean diet adherence and disease risk, Pearson correlations were calculated between food scores (MED-Diet adherence) and each individual disease risk score, as well as the cumulative disease risk score, across all participants and timepoints (baseline and follow-up, both study arms). P-values were adjusted for multiple comparisons using the Benjamini-Hochberg false discovery rate (FDR) method, and associations were considered significant at FDR $\leq 0.05$. To assess longitudinal relationships, change scores (Δ) were calculated for each disease risk score and for clinical/metadata variables as the difference between follow-up and baseline values within participants with paired measurements at both timepoints, across both study arms. Pearson correlations were computed between Δ disease risk scores and Δ values for constructional praxis score and circulating IL-17 levels. Associations were considered nominally significant at $p \leq 0.05$. All correlation analyses were performed using the corr.test function from the 'psych' package in R, with pairwise complete observations used for the longitudinal (Δ) analyses. Scatter plots of effect size (Pearson's r) against $-\log_{10}$(adjusted p-value) or $-\log_{10}$(nominal p-value) were generated using ggplot2, with point labels added using ggrepel.

To determine whether MifRix-derived disease risk-scores reflect therapeutic response, changes in Generic Disease (GD) risk-scores before and after fecal microbiota transplantation (FMT) were evaluated in three independent intervention studies (DavarD_2023, KongL_2020 and KumpP_2017). For each participant, the change in predicted disease risk (Δ risk-score = post − pre) was calculated and compared between responders and non-responders using two-sided Wilcoxon rank-sum tests. Differences with $p \leq 0.05$ were considered statistically significant.

The relationship between predicted disease risk and clinical improvement following FMT was further investigated using an independent Indian cohort (PRJEB115184) comprising 75 patients with Crohn's disease (CD). MifRix-derived GD risk-scores were first compared between healthy stool donors and CD patients using two-sided Wilcoxon rank-sum tests to assess whether predicted disease risk distinguished healthy and diseased microbiomes. Changes in GD risk-scores following FMT (Δ risk-score = post − pre) were subsequently compared with changes in Crohn's Disease Activity Index (CDAI) scores. Patients were stratified according to the magnitude of risk-score reduction, and differences in CDAI changes among groups were assessed using Dunn's multiple comparison test following Kruskal-Wallis analysis. Finally, changes in GD risk-scores were compared between patients achieving clinical remission and non-remitters using two-sided Wilcoxon rank-sum tests to determine whether reductions in predicted disease risk were associated with treatment response.

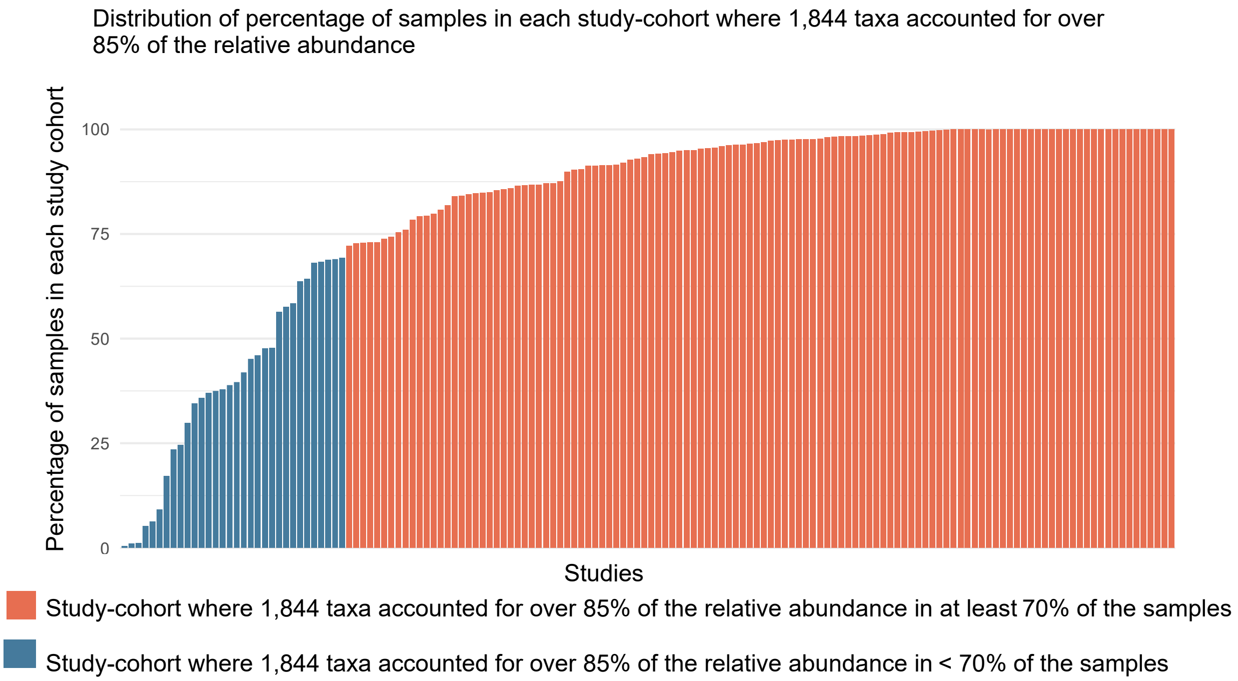


**Fig. S1.** Bar plots showing the percentage of microbiomes in each study-cohorts shown on the X-axis where the 1,844 taxa showed a relative abundance >= 70%. The orange-coloured bars show the study-cohorts where 1,844 taxa accounted for over 85% of the relative abundance in at least 70% of the samples in each cohort, on the other hand the blue-coloured bars show study-cohorts where 1,844 taxa accounted for over 85% of the relative abundance in less than 70% of the samples. These 1,844 taxa accounted for over 85% of relative abundance in overall 87% samples irrespective of specific study-cohorts.

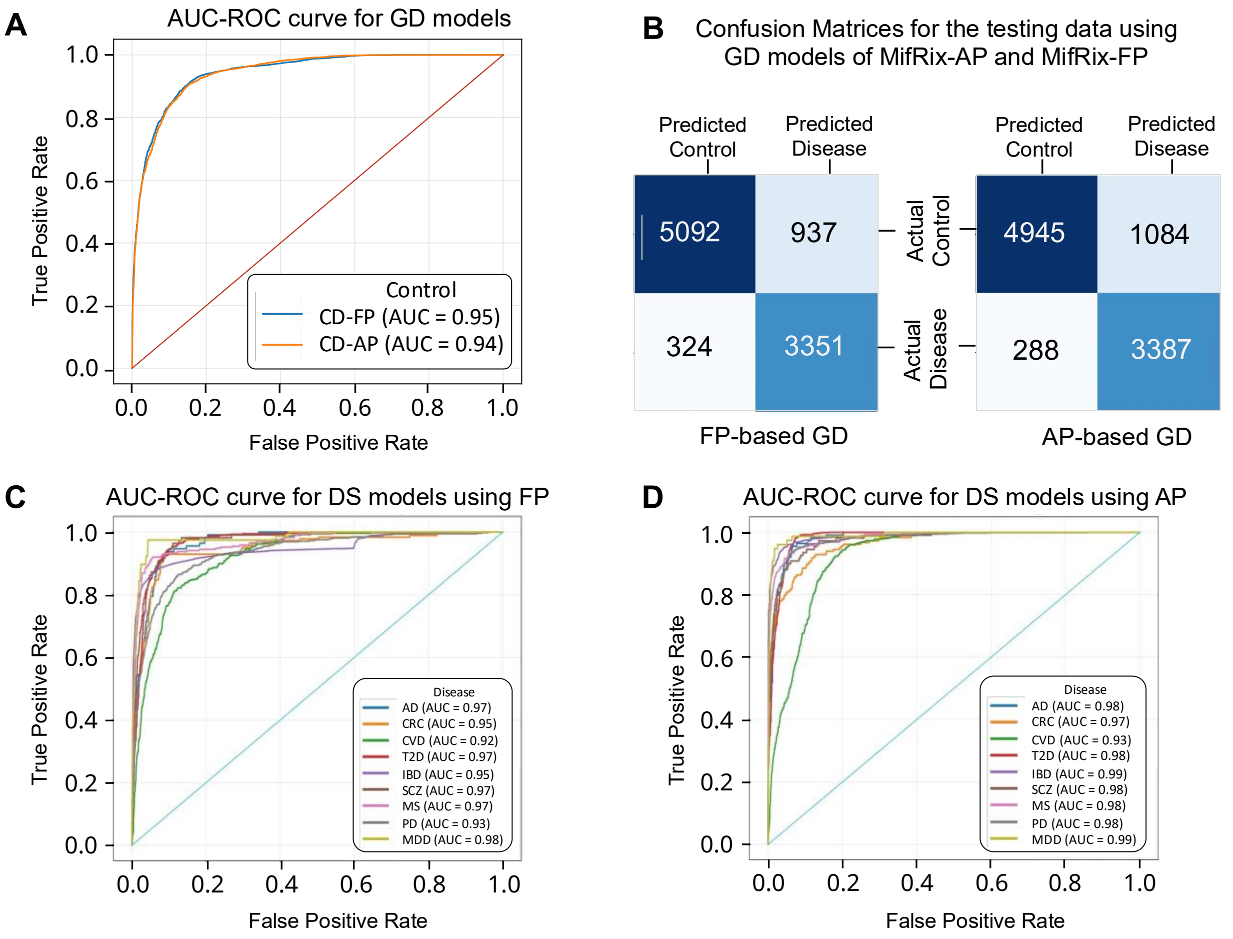


**Fig. S2. A.** AUC-ROC curves of MifRix 'Generic-Disease' (GD) models (both MifRix-AP and MifRix-FP variants) on the testing subset of the training sub-cohorts. **B.** Confusion Matrices for evaluating the classification performances of the GD models of the MifRix-AP variant (right) and MifRix-FP variant (left). **A** & **B** show that both variants of the GD model show similar performance for the testing subset of the training study-cohorts. **C.** AUC-ROC curves of the 'Disease-Specific' (DS) models for the MifRix-FP variant for the 10 diseases. **D.** AUC-ROC curves of the 'Disease-Specific' (DS) models for the MifRix-AP variant for the 10 diseases. Both **C** & **D** show the results for specifically the testing subset microbiomes of the training study-cohorts.

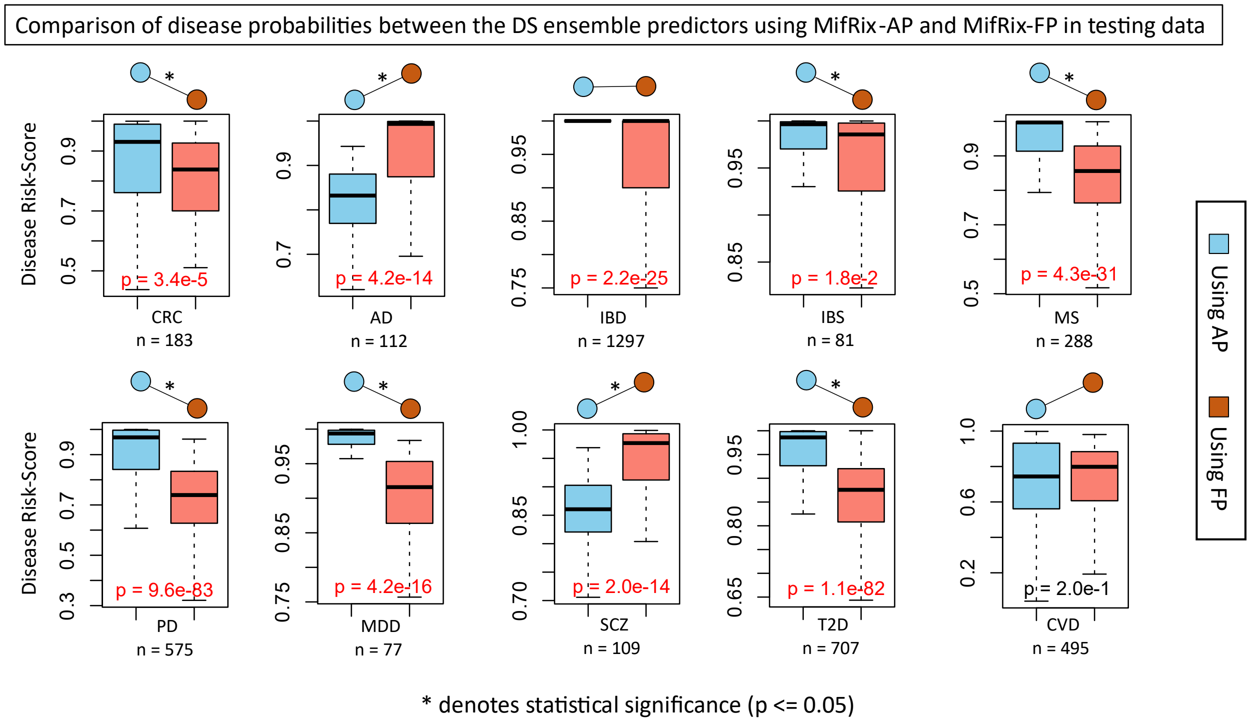


**Fig. S3.** Comparing the disease risk-scores of microbiomes from particular diseases in the testing dataset using the DS (Disease-specific) ensemble predictors of MifRix-AP and MifRix-FP. In 6 out of 10 diseases, the disease risk-scores predicted by MifRix-AP were significantly higher (p-value < 0.05; using Wilcoxon signed-rank test) as compared to risks generated using MifRix-FP whereas in two diseases MifRix-FP were significantly higher than MifRix-AP. The two bulbs joined through a line shows the pattern (increase/decrease/equal) of the medians across the boxplots and the '*' denotes the significance level (p-value <= 0.05).

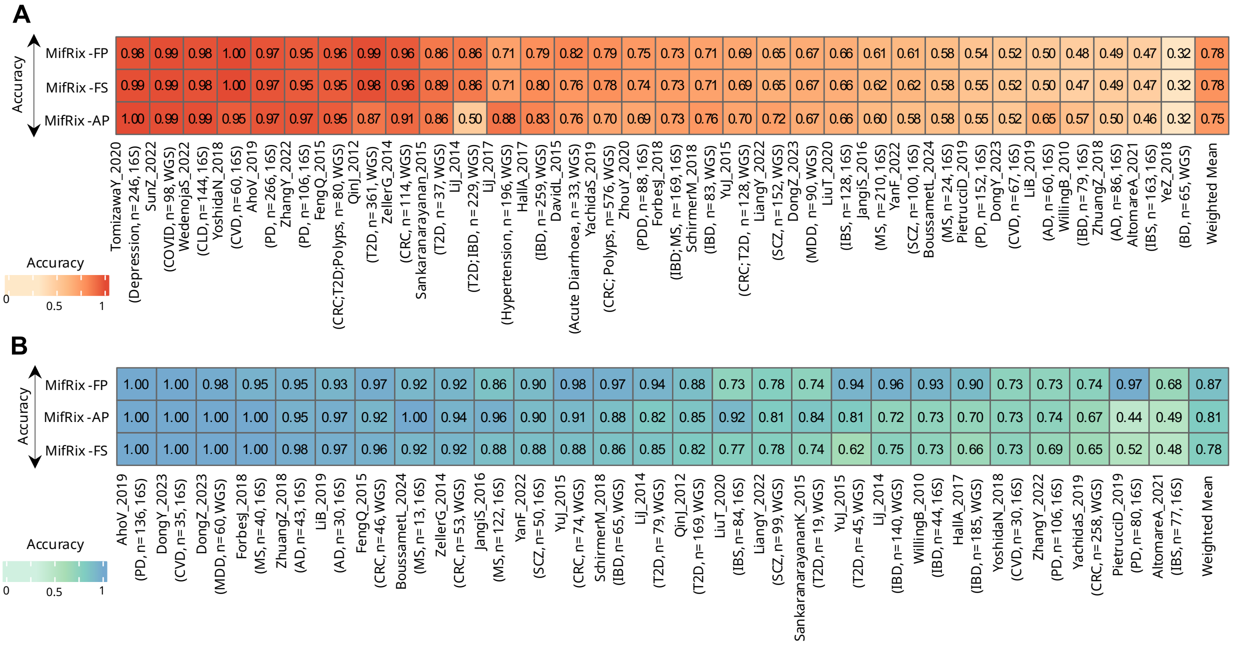

Note: MifRix -FS denotes MifRix -final-score

**Fig. S4. A.** Heatmap comparing classification accuracy of the generic disease (GD) risk-scores from the three MifRix variants (MifRix-AP, MifRix-FP, MifRix-final-score) across the 32 unseen validation cohorts. Colour scale reflects accuracy (0-1), with exact values annotated in each cell; the rightmost column shows the weighted mean accuracy across all cohorts for MifRix-FP (78%), MifRix-final-score (78%), and MifRix-AP (75%). **B.** Heatmap comparing classification accuracy of the disease-specific (DS) risk-scores from the same three variants across the 27 unseen validation cohorts, formatted as in (A); weighted mean accuracy was 87% (MifRix-FP), 81% (MifRix-AP), and 78% (MifRix-final-score). For both panels, classification accuracy was calculated by predicting each microbiome as diseased (GD) or as belonging to the target disease (DS) when the corresponding risk-score was > 0.5, and comparing these predictions against the true disease labels.

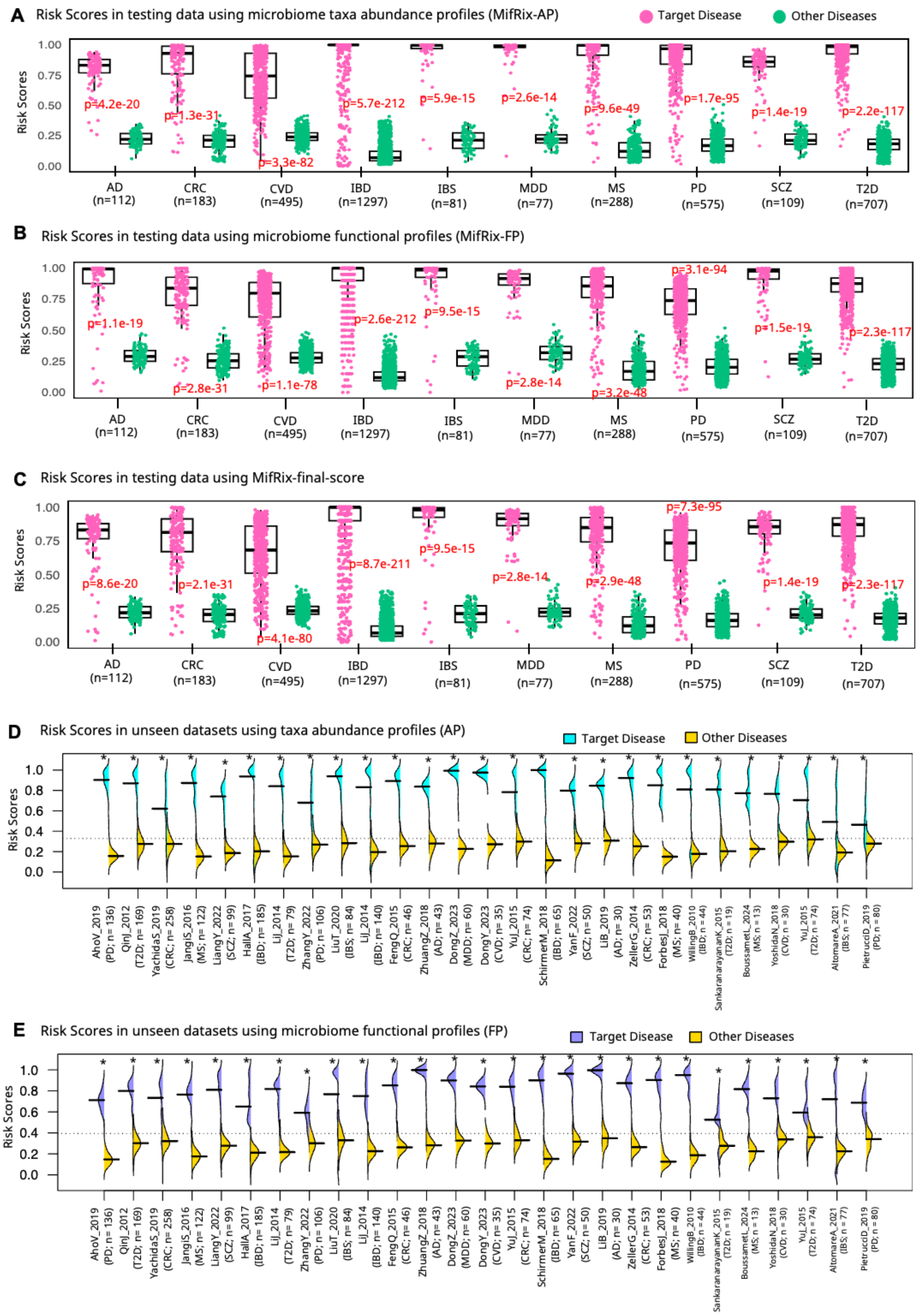

A Risk Scores in testing data using microbiome taxa abundance profiles (MifRix-AP)
Target Disease
Other Diseases
Risk Scores
1.00
0.75
0.50
0.25
0.00
p=4.2e-20
p=1.3e-31
p=3.3e-82
p=5.7e-212
p=5.9e-15
p=2.6e-14
p=9.6e-49
p=1.7e-95
p=1.4e-19
p=2.2e-117
AD (n=112)
CRC (n=183)
CVD (n=495)
IBD (n=1297)
IBS (n=81)
MDD (n=77)
MS (n=288)
PD (n=575)
SCZ (n=109)
T2D (n=707)
B Risk Scores in testing data using microbiome functional profiles (MifRix-FP)
p=1.1e-19
p=2.8e-31
p=1.1e-78
p=2.6e-212
p=9.5e-15
p=2.8e-14
p=3.2e-48
p=3.1e-94
p=1.5e-19
p=2.3e-117
C Risk Scores in testing data using MifRix-final-score
p=8.6e-20
p=2.1e-31
p=4.1e-80
p=8.7e-211
p=9.5e-15
p=2.8e-14
p=2.9e-48
p=7.3e-95
p=1.4e-19
p=2.3e-117
D Risk Scores in unseen datasets using taxa abundance profiles (AP)
E Risk Scores in unseen datasets using microbiome functional profiles (FP)
1.0
0.8
0.6
0.4
0.2
0.0
AhoV_2019 (PD; n= 136)
QinJ_2012 (T2D; n= 169)
YachidaS_2019 (CRC; n= 258)
JangIS_2016 (MS; n= 122)
LiangY_2022 (SCZ; n= 99)
HallA_2017 (IBD; n= 185)
LiJ_2014 (T2D; n= 79)
ZhangY_2022 (PD; n= 106)
LiuT_2020 (IBS; n= 84)
LiJ_2014 (IBD; n= 140)
FengQ_2015 (CRC; n= 46)
ZhuangZ_2018 (AD; n= 43)
DongZ_2023 (MDD; n= 60)
DongY_2023 (CVD; n= 35)
YuJ_2015 (CRC; n= 74)
SchirmerM_2018 (IBD; n= 65)
YanF_2022 (SCZ; n= 50)
LiB_2019 (AD; n= 30)
ZellerG_2014 (CRC; n= 53)
ForbesJ_2018 (MS; n= 40)
WillingB_2010 (IBD; n= 44)
SankaranarayananK_2015 (T2D; n = 19)
BoussametL_2024 (MS; n = 13)
YoshidaN_2018 (CVD; n = 30)
YuJ_2015 (T2D; n = 74)
AltomareA_2021 (IBS; n = 77)
PietrucciD_2019 (PD; n = 80)

**Fig. S5. MiFRix Disease-specific model generated risk scores can distinguish specific disease from other diseases using both microbiome taxa abundance profiles as well as microbiome functional profiles.** Boxplot comparing the distributions of the DS-risk-scores for each of the target diseases (indicated on the X-axis) with the mean DS-risk-scores of the corresponding nine non-target diseases for each of the scenarios, when the DS-risk-scores are generated using **A.** MifRix-AP **B.** MifRix-FP and **C.** MifRix-final-score variants**.** In **A-C,** the comparisons are performed for the microbiomes constituting the testing subset of the training cohorts. Each disease is annotated with the corresponding disease name and sample size (n) on the X-axis label. Statistical significance between the two distributions was assessed using the Wilcoxon signed-rank test; *p*-values shown in red indicate significant differences ($p < 0.05$). The same comparisons of the 27 unseen validation cohorts are provided in **D-E** (comparisons being shown as bean plots), where **D.** provides the comparison of the DS-risk-scores generated by the MifRix-AP and **E.** provides the comparison of the DS-risk-scores generated by MifRix-FP. The same comparisons within the validation cohort microbiomes using the MifRix-final-score are provided in **Fig. 3C**. Each bean plot is annotated with the corresponding disease name, study cohort and sample size (n) on the X-axis label. Statistical significance between the two distributions was assessed using the Wilcoxon signed-rank test; *p*-values shown in red indicate significant differences ($p < 0.05$).

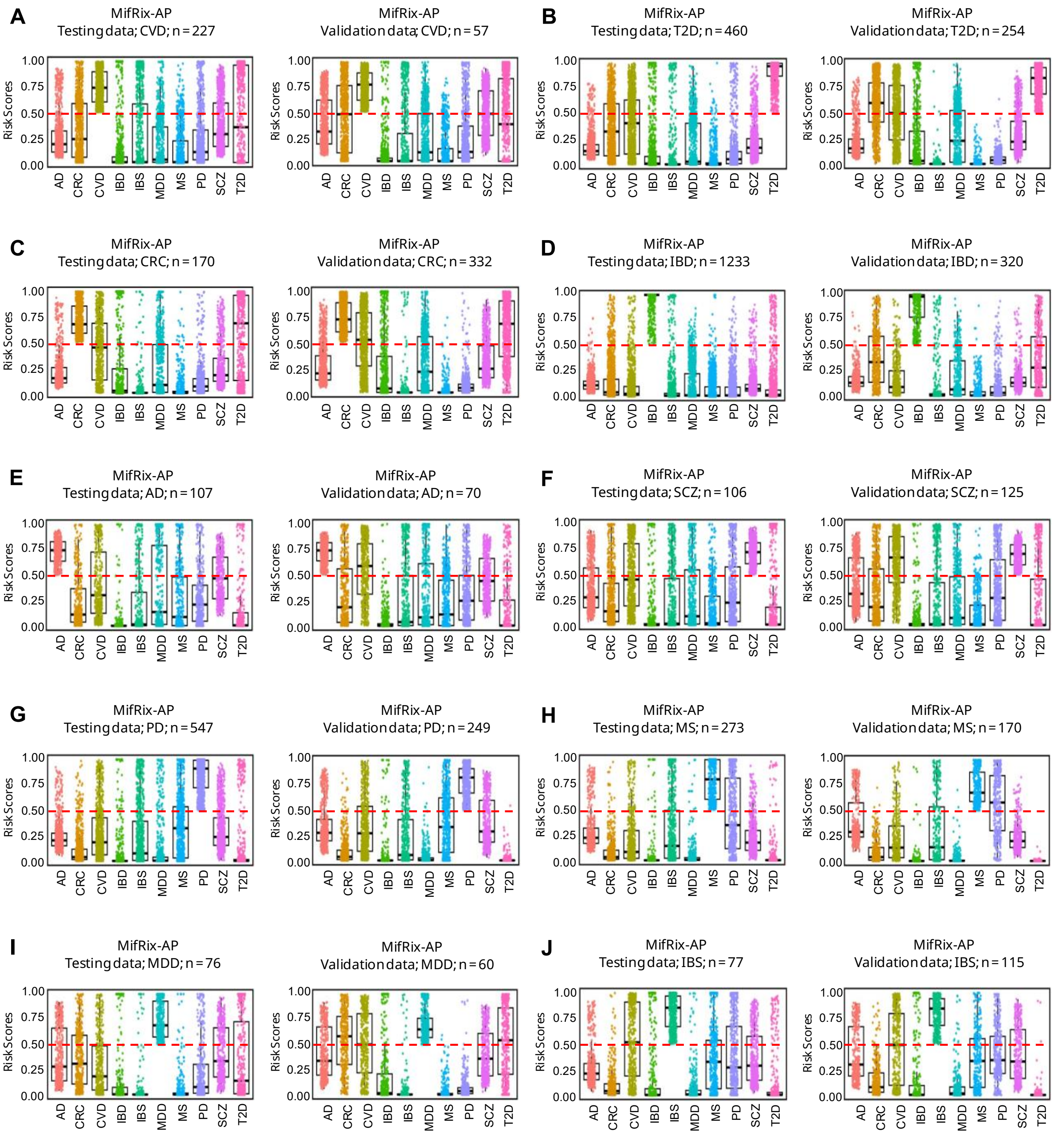


**Fig. S6. A-J.** Boxplots comparing the distribution of MifRix-AP-derived disease-specific risk-scores across ten diseases for microbiomes predicted as high-risk (MifRix-AP >0.5) for the corresponding diseases, where the target diseases are CVD (**A**) T2D (**B**), CRC (**C**), IBD (**D**), AD (**E**), SCZ (**F**), PD (**G**), MS (**H**), MDD (**I**) and IBS (**J**). In each panel (**A-J**), the comparative boxplots were generated both the testing subset of the training sub-cohorts (left) as well as unseen validation datasets (right). Each boxplot was overlaid with jittered data points (each data point indicates a microbiome). These results for the MifRix-final-score are shown in **Figs. 3D-G** and **4A-F**.

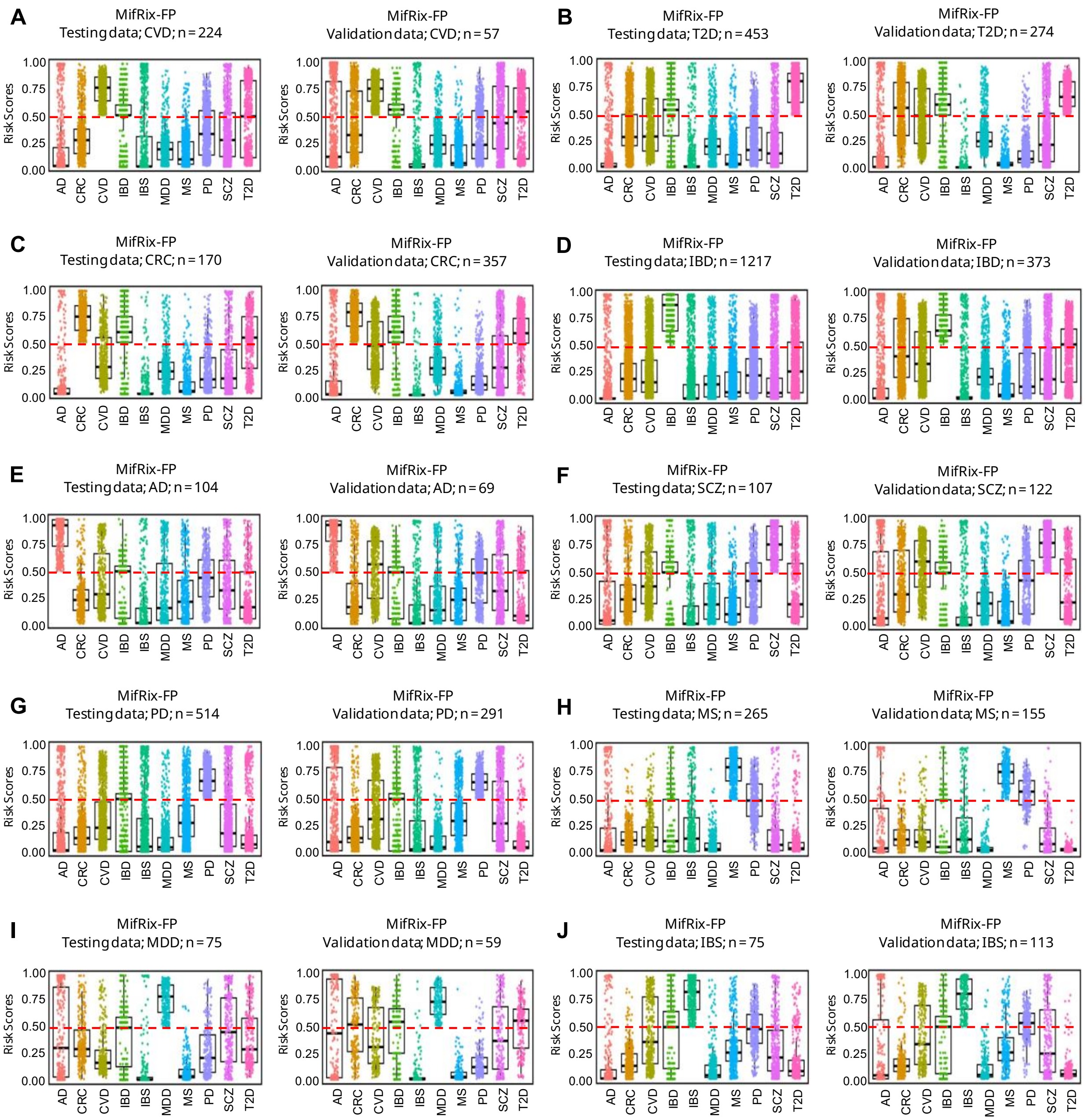


**Fig. S7. A-J.** Boxplots comparing the distribution of MifRix-FP-derived disease-specific risk-scores across ten diseases for microbiomes predicted as high-risk (MifRix-FP >0.5) for the corresponding diseases, where the target diseases are CVD (**A**) T2D (**B**), CRC (**C**), IBD (**D**), AD (**E**), SCZ (**F**), PD (**G**), MS (**H**), MDD (**I**) and IBS (**J**). In each panel (**A-J**), the comparative boxplots were generated both the testing subset of the training sub-cohorts (left) as well as unseen validation datasets (right). Each boxplot was overlaid with jittered data points (each data point indicates a microbiome). These results for the MifRix-final-score are shown in **Figs. 3D-G** and **4A-F**.

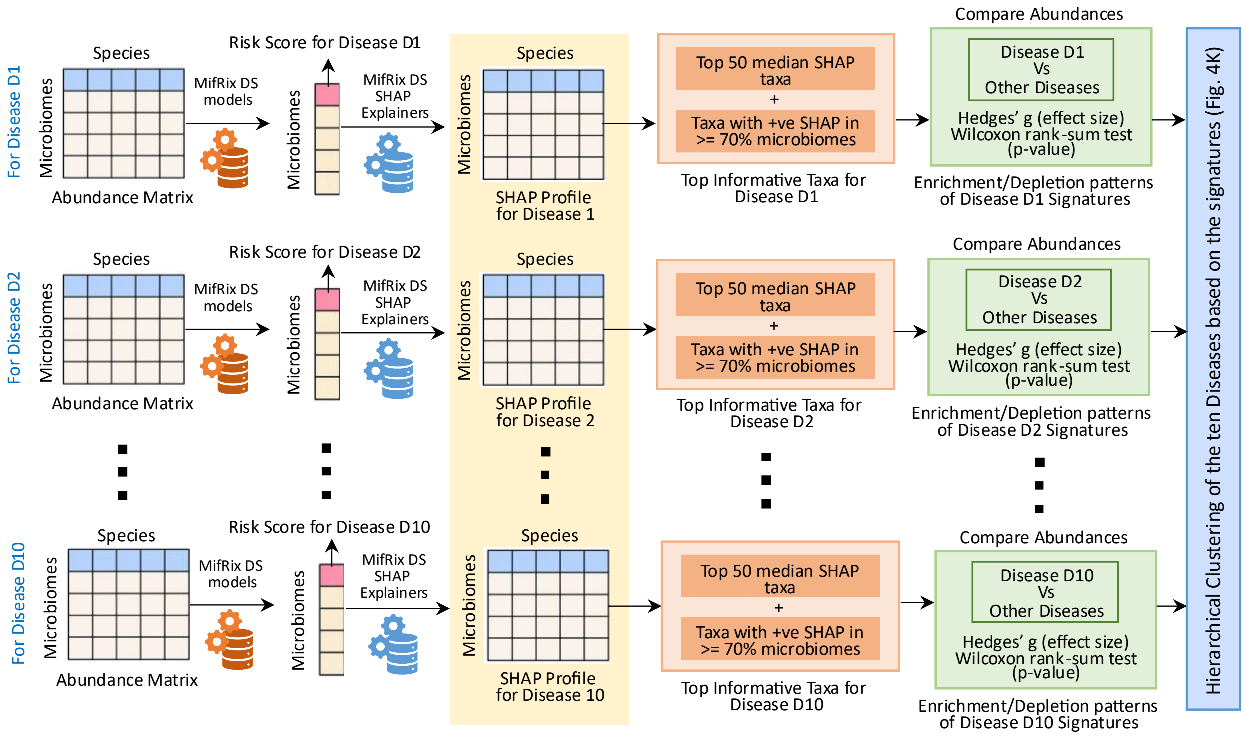


**Fig. S8. Workflow for deriving disease-specific microbial signatures from MifRix-AP risk prediction models.** Disease-specific microbiome profiles are first analyzed using MifRix-AP ensemble models to generate disease risk scores. Model predictions are subsequently interpreted using SHapley Additive exPlanations (SHAP), with SHAP values computed using TreeExplainer or LinearExplainer according to the underlying model type and aggregated across ensemble models using the median. For each disease, microbial taxa are ranked by their median SHAP values, and the most informative taxa are selected based on positive feature contributions and consistent occurrence (positive SHAP value in at least 70% of the microbiomes) across microbiomes. The abundances of the selected taxa are then compared between microbiomes from the target disease and those from all other diseases using two-sided Wilcoxon rank-sum tests, with effect sizes quantified using Hedges' g. Taxa are finally classified as enriched, depleted, or not significantly altered, generating a disease-specific directionality matrix that summarizes microbial signature patterns across diseases.

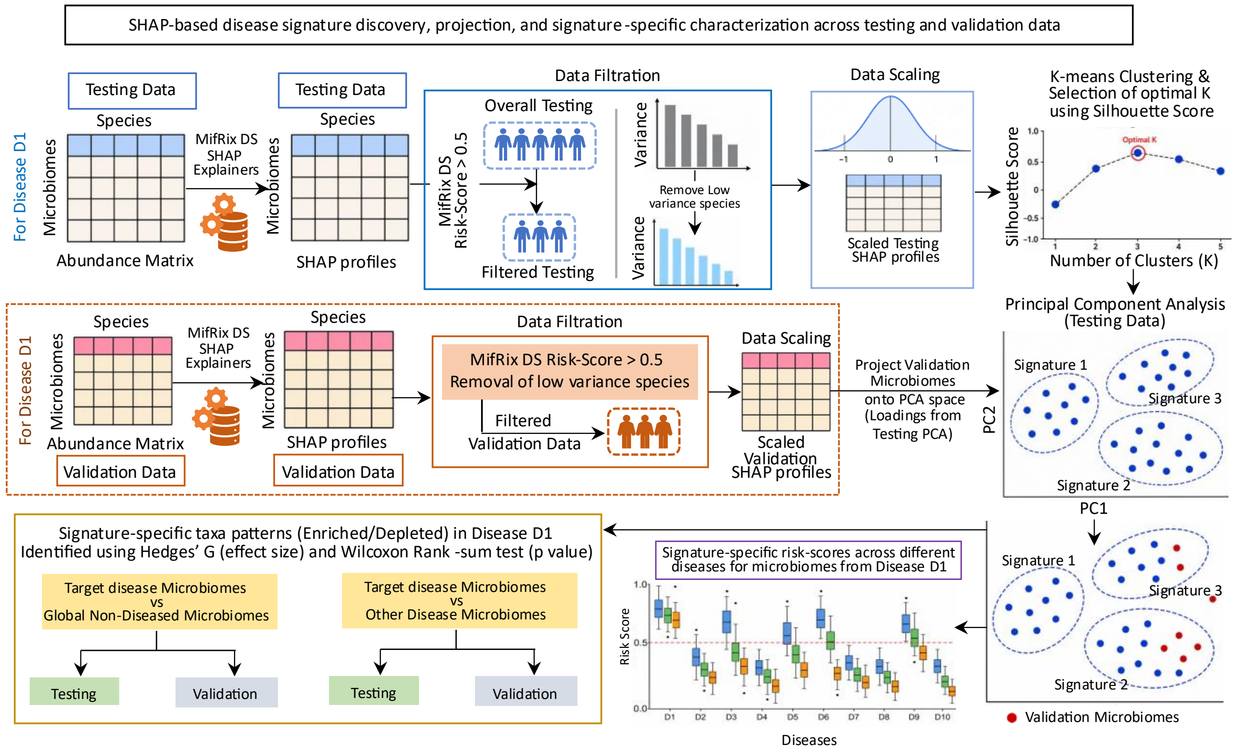


**Fig. S9. Identification of distinct, reproducible sub-signatures of microbiome alteration within the same disease using SHAP-based signature discovery.** Schematic of the workflow used for SHAP-based signature discovery, projection, and cluster-specific characterization across testing and validation cohorts. Disease-specific MifRix ensemble models were applied to microbiome abundance profiles to generate species-level SHAP value matrices, computed independently for the testing subset of the training cohorts and for unseen validation cohorts. Within the testing subset, disease-associated microbiomes (target-disease risk-score >0.5) were retained, low-variance features removed, and SHAP values z-scaled. K-means clustering was performed on the resulting matrix, with the optimal cluster number determined via silhouette analysis; each resulting cluster represented a distinct microbiome sub-signature for the same target disease. Principal component analysis (PCA) was then performed on the same scaled SHAP matrix, with testing microbiomes visualized according to their cluster (sub-signature) assignment. Each microbiome in the unseen validation datasets for a given was projected into this testing-subset-derived PCA space using the corresponding scaling parameters and PCA loadings, and was assigned to the sub-signature cluster whose centroid was closest to it. Risk prediction scores for all ten disease models were compared across microbiome signatures in both testing and validation cohorts to assess disease-specific and cross-disease risk distributions. Finally, cluster-specific microbial signatures were identified by comparing microbiomes within each signature against (i) global healthy controls and (ii) microbiomes from all other diseases separately, with the direction and magnitude of enrichment or depletion quantified using Hedges' *g* and statistical significance assessed using the Wilcoxon rank-sum test.

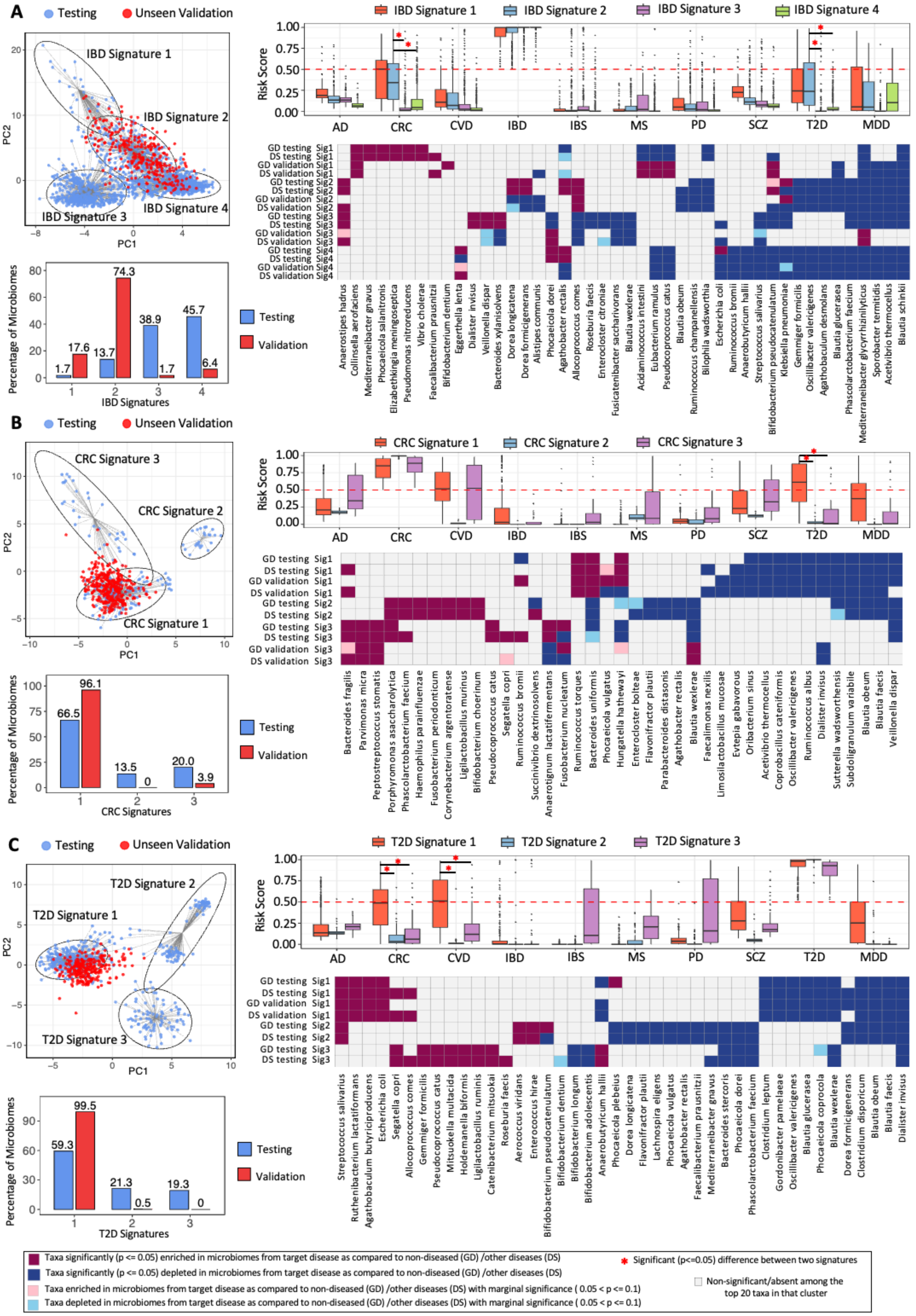
A
Testing
Unseen Validation
IBD Signature 1
IBD Signature 2
IBD Signature 3
IBD Signature 4
PC1
PC2
Percentage of Microbiomes
IBD Signatures
Testing
Validation
IBD Signature 1
IBD Signature 2
IBD Signature 3
IBD Signature 4
Risk Score
AD
CRC
CVD
IBD
IBS
MS
PD
SCZ
T2D
MDD
GD testing Sig1
DS testing Sig1
GD validation Sig1
DS validation Sig1
GD testing Sig2
DS testing Sig2
GD validation Sig2
DS validation Sig2
GD testing Sig3
DS testing Sig3
GD validation Sig3
DS validation Sig3
GD testing Sig4
DS testing Sig4
GD validation Sig4
DS validation Sig4
Anaerostipes hadrus
Collinsella aerofaciens
Mediterraneibacter gnavus
Phocaeicola salanitronis
Elizabethkingia meningoseptica
Pseudomonas nitroreducens
Vibrio cholerae
Faecalibacterium prausnitzii
Bifidobacterium dentium
Eggerthella lenta
Dialister invisus
Veillonella dispar
Bacteroides xylanisolvens
Dorea longicatena
Dorea formicigenerans
Alistipes communis
Phocaeicola dorei
Agathobacter rectalis
Allocoprococcus comes
Roseburia faecis
Enterocloster citroniae
Fusicatenibacter saccharivorans
Blautia wexlerae
Acidaminococcus intestini
Eubacterium ramulus
Pseudocoprococcus catus
Blautia obeum
Ruminococcus champanellensis
Bilophila wadsworthia
Escherichia coli
Ruminococcus bromii
Anaerobutyricum hallii
Streptococcus salivarius
Bifidobacterium pseudocatenulatum
Klebsiella pneumoniae
Gemmiger formicilis
Oscillibacter valericigenes
Agathobaculum desmolans
Blautia glucerasea
Phascolarctobacterium faecium
Mediterraneibacter glycyrrhizinilyticus
Sporobacter termitidis
Acetivibrio thermocellus
Blautia schinkii
B
Testing
Unseen Validation
CRC Signature 3
CRC Signature 2
CRC Signature 1
PC1
PC2
Percentage of Microbiomes
CRC Signatures
Testing
Validation
CRC Signature 1
CRC Signature 2
CRC Signature 3
Risk Score
AD
CRC
CVD
IBD
IBS
MS
PD
SCZ
T2D
MDD
GD testing Sig1
DS testing Sig1
GD validation Sig1
DS validation Sig1
GD testing Sig2
DS testing Sig2
GD testing Sig3
DS testing Sig3
GD validation Sig3
DS validation Sig3
Bacteroides fragilis
Parvimonas micra
Peptostreptococcus stomatis
Porphyromonas asaccharolytica
Phascolarctobacterium faecium
Haemophilus parainfluenzae
Fusobacterium periodonticum
Corynebacterium argentoratense
Ligilactobacillus murinus
Bifidobacterium choerinum
Pseudocoprococcus catus
Segatella copri
Ruminococcus bromii
Succinivibrio dextrinosolvens
Anaerotignum lactatifermentans
Fusobacterium nucleatum
Ruminococcus torques
Bacteroides uniformis
Phocaeicola vulgatus
Hungatella hathewayi
Enterocloster bolteae
Flavonifractor plautii
Parabacteroides distasonis
Agathobacter rectalis
Blautia wexlerae
Faecalimonas nexilis
Limosilactobacillus mucosae
Evtepia gabavorous
Oribacterium sinus
Acetivibrio thermocellus
Coprobacillus cateniformis
Oscillibacter valericigenes
Ruminococcus albus
Dialister invisus
Sutterella wadsworthensis
Subdoligranulum variabile
Blautia obeum
Blautia faecis
Veillonella dispar
C
Testing
Unseen Validation
T2D Signature 1
T2D Signature 2
T2D Signature 3
PC1
PC2
Percentage of Microbiomes
T2D Signatures
Testing
Validation
T2D Signature 1
T2D Signature 2
T2D Signature 3
Risk Score
AD
CRC
CVD
IBD
IBS
MS
PD
SCZ
T2D
MDD
GD testing Sig1
DS testing Sig1
GD validation Sig1
DS validation Sig1
GD testing Sig2
DS testing Sig2
GD testing Sig3
DS testing Sig3
Streptococcus salivarius
Ruthenibacterium lactatiformans
Agathobaculum butyriciproducens
Escherichia coli
Segatella copri
Allocoprococcus comes
Gemmiger formicilis
Pseudocoprococcus catus
Mitsuokella multacida
Holdemanella biformis
Ligilactobacillus ruminis
Catenibacterium mitsuokai
Roseburia faecis
Aerococcus viridans
Enterococcus hirae
Bifidobacterium pseudocatenulatum
Bifidobacterium dentium
Bifidobacterium longum
Bifidobacterium adolescentis
Anaerobutyricum hallii
Phocaeicola plebeius
Dorea longicatena
Flavonifractor plautii
Lachnospira eligens
Phocaeicola vulgatus
Agathobacter rectalis
Faecalibacterium prausnitzii
Mediterraneibacter gnavus
Bacteroides stercoris
Phocaeicola dorei
Phascolarctobacterium faecium
Clostridium leptum
Gordonibacter pamelaeae
Oscillibacter valericigenes
Blautia glucerasea
Phocaeicola coprocola
Blautia wexlerae
Dorea formicigenerans
Clostridium disporicum
Blautia obeum
Blautia faecis
Dialister invisus
Taxa significantly (p <= 0.05) enriched in microbiomes from target disease as compared to non-diseased (GD) /other diseases (DS)
Taxa significantly (p <= 0.05) depleted in microbiomes from target disease as compared to non-diseased (GD) /other diseases (DS)
Taxa enriched in microbiomes from target disease as compared to non-diseased (GD) /other diseases (DS) with marginal significance ( 0.05 < p <= 0.1)
Taxa depleted in microbiomes from target disease as compared to non-diseased (GD) /other diseases (DS) with marginal significance ( 0.05 < p <= 0.1)
* Significant (p<=0.05) difference between two signatures
Non-significant/absent among the top 20 taxa in that cluster

**Fig. S10. Disease-specific microbiome signatures identified from explainable AI-derived SHAP landscapes are reproducible across independent validation cohorts. A-C**. Disease-specific microbiome signatures identified for Inflammatory Bowel Disorder (IBD) (**A**), Colorectal Cancer (CRC) (**B**), and type 2 diabetes (T2D) (**C**) based on clustering of microbiome SHAP profiles generated using the corresponding disease-specific MifRix ensemble models. For each disease, the left panel shows principal component analysis (PCA) of SHAP values from the testing cohort (as blue points), with microbiome signatures defined by k-means clustering. Microbiomes from the independent validation cohort (shown as red points) were projected onto the PCA space derived from the testing cohort to assess signature reproducibility. The upper middle panel of box plot compares predicted risk scores for ten diseases (Alzheimer's Disease (AD), CRC, Cardiovascular Diseases (CVD), IBD, Irritable Bowel Syndrome (IBS), Multiple Sclerosis (MS), Parkinson's Disease (PD), Schizophrenia (SCZ), T2D and Major Depressive Disorder (MDD)) across the identified microbiome signatures. Red dashed lines indicate the disease-specific risk threshold (which is 0.5), and asterisks denote significant differences between signatures ($p \leq 0.05$) using Wilcoxon rank-sum test. The lower middle panel of bar plot shows the proportion of testing and validation microbiomes assigned to each signature. Right panels of heatmaps summarize signature-associated microbial taxa that were enriched or depleted relative to microbiomes from the target disease versus non-diseased controls (GD) or other diseases (DS) in both testing and validation datasets. Taxa are classified according to Hedges' *g* effect size and Wilcoxon rank-sum test significance. Heatmap cells are colored according to the direction and statistical significance of differential abundance relative to microbiomes from the target disease versus non-diseased controls (GD) or other diseases (DS) across both testing and validation data. Dark magenta indicates significant enrichment (Hedges' $g > 0$ & $p \leq 0.05$); dark blue indicates significant depletion (Hedges' $g < 0$ & $p \leq 0.05$); light pink indicates marginal enrichment (Hedges' $g > 0$ & $0.05 < p \leq 0.1$); light blue indicates marginal depletion (Hedges' $g < 0$ & $0.05 < p \leq 0.1$); and white denotes non-significant or absent among the top 20 taxa for that signature.

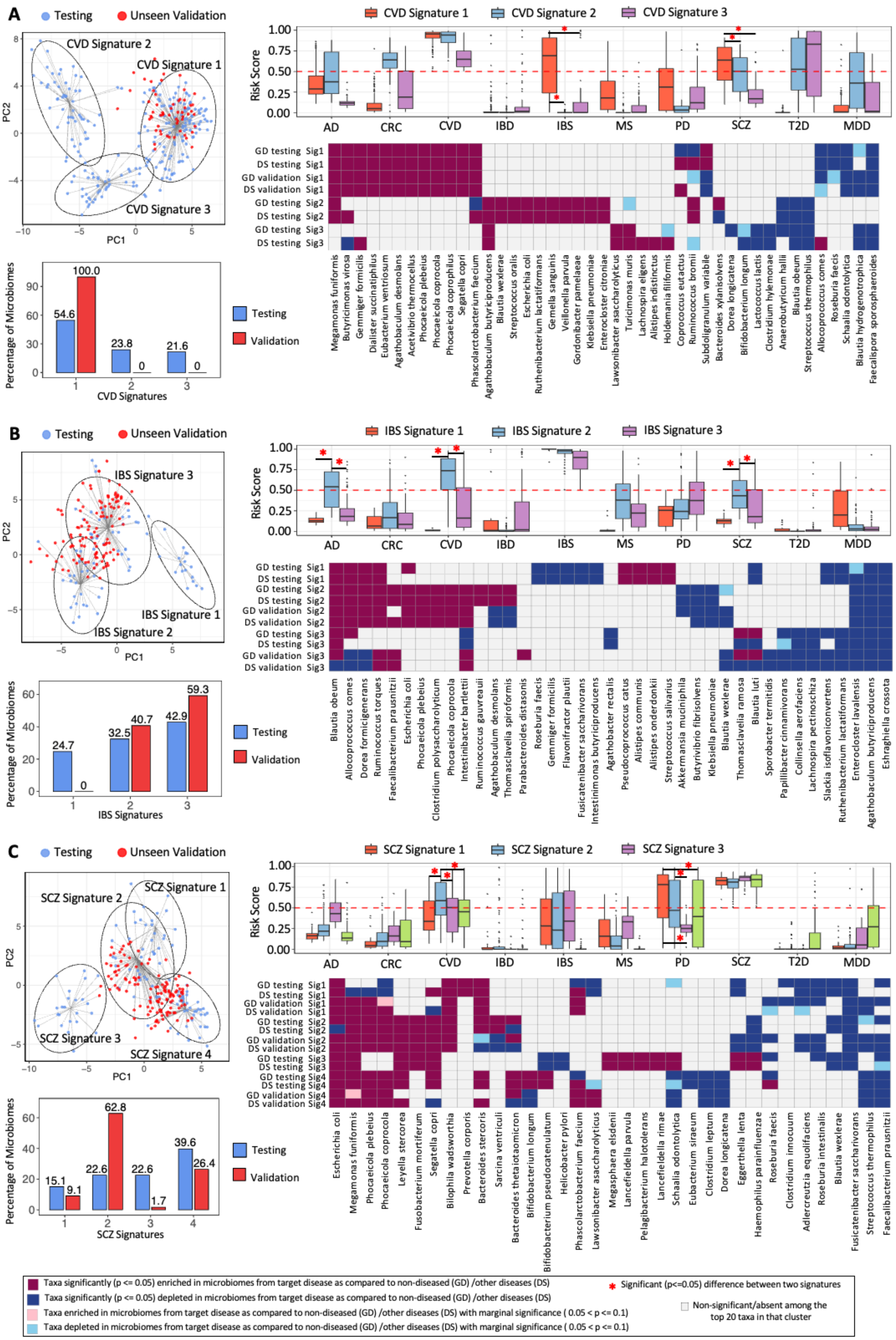
A
Testing
Unseen Validation
CVD Signature 1
CVD Signature 2
CVD Signature 3
PC1
PC2
Percentage of Microbiomes
54.6
100.0
23.8
0
21.6
0
CVD Signatures
Testing
Validation
Risk Score
AD
CRC
CVD
IBD
IBS
MS
PD
SCZ
T2D
MDD
GD testing Sig1
DS testing Sig1
GD validation Sig1
DS validation Sig1
GD testing Sig2
DS testing Sig2
GD testing Sig3
DS testing Sig3
Megamonas funiformis
Butyricimonas virosa
Gemmiger formicilis
Dialister succinatiphilus
Eubacterium ventriosum
Agathobaculum desmolans
Acetivibrio thermocellus
Phocaeicola plebeius
Phocaeicola coprocola
Phocaeicola coprophilus
Segatella copri
Phascolarctobacterium faecium
Agathobaculum butyriciproducens
Blautia wexlerae
Streptococcus oralis
Escherichia coli
Ruthenibacterium lactatiformans
Gemella sanguinis
Veillonella parvula
Gordonibacter pamelaeae
Klebsiella pneumoniae
Enterocloster citroniae
Lawsonibacter asaccharolyticus
Turicimonas muris
Lachnospira eligens
Alistipes indistinctus
Holdemania filiformis
Coprococcus eutactus
Ruminococcus bromii
Subdoligranulum variabile
Bacteroides xylanisolvens
Dorea longicatena
Bifidobacterium longum
Lactococcus lactis
Clostridium hylemonae
Anaerobutyricum hallii
Blautia obeum
Streptococcus thermophilus
Allocoprococcus comes
Roseburia faecis
Schaalia odontolytica
Blautia hydrogenotrophica
Faecalispora sporosphaeroides
B
Testing
Unseen Validation
IBS Signature 1
IBS Signature 2
IBS Signature 3
PC1
PC2
Percentage of Microbiomes
24.7
0
32.5
40.7
42.9
59.3
IBS Signatures
Testing
Validation
Risk Score
AD
CRC
CVD
IBD
IBS
MS
PD
SCZ
T2D
MDD
GD testing Sig1
DS testing Sig1
GD testing Sig2
DS testing Sig2
GD validation Sig2
DS validation Sig2
GD testing Sig3
DS testing Sig3
GD validation Sig3
DS validation Sig3
Blautia obeum
Allocoprococcus comes
Dorea formicigenerans
Ruminococcus torques
Faecalibacterium prausnitzii
Escherichia coli
Phocaeicola plebeius
Clostridium polysaccharolyticum
Phocaeicola coprocola
Intestinibacter bartlettii
Ruminococcus gauvreauii
Agathobaculum desmolans
Thomasclavelia spiroformis
Parabacteroides distasonis
Roseburia faecis
Gemmiger formicilis
Flavonifractor plautii
Fusicatenibacter saccharivorans
Intestinimonas butyriciproducens
Agathobacter rectalis
Pseudocoprococcus catus
Alistipes communis
Alistipes onderdonkii
Streptococcus salivarius
Akkermansia muciniphila
Butyrivibrio fibrisolvens
Klebsiella pneumoniae
Blautia wexlerae
Thomasclavelia ramosa
Blautia luti
Sporobacter termitidis
Papillibacter cinnamivorans
Collinsella aerofaciens
Lachnospira pectinoschiza
Slackia isoflavoniconvertens
Ruthenibacterium lactatiformans
Enterocloster lavalensis
Agathobaculum butyriciproducens
Eshraghiella crossota
C
Testing
Unseen Validation
SCZ Signature 1
SCZ Signature 2
SCZ Signature 3
SCZ Signature 4
PC1
PC2
Percentage of Microbiomes
15.1
9.1
22.6
62.8
22.6
1.7
39.6
26.4
SCZ Signatures
Testing
Validation
Risk Score
AD
CRC
CVD
IBD
IBS
MS
PD
SCZ
T2D
MDD
GD testing Sig1
DS testing Sig1
GD validation Sig1
DS validation Sig1
GD testing Sig2
DS testing Sig2
GD validation Sig2
DS validation Sig2
GD testing Sig3
DS testing Sig3
GD testing Sig4
DS testing Sig4
GD validation Sig4
DS validation Sig4
Escherichia coli
Megamonas funiformis
Phocaeicola plebeius
Phocaeicola coprocola
Leyella stercorea
Fusobacterium mortiferum
Segatella copri
Bilophila wadsworthia
Prevotella corporis
Bacteroides stercoris
Sarcina ventriculi
Bacteroides thetaiotaomicron
Bifidobacterium longum
Bifidobacterium pseudocatenulatum
Helicobacter pylori
Phascolarctobacterium faecium
Lawsonibacter asaccharolyticus
Megasphaera elsdenii
Lancefieldella parvula
Pelagibacterium halotolerans
Lancefieldella rimae
Schaalia odontolytica
Eubacterium siraeum
Clostridium leptum
Dorea longicatena
Eggerthella lenta
Haemophilus parainfluenzae
Roseburia faecis
Clostridium innocuum
Adlercreutzia equolifaciens
Roseburia intestinalis
Blautia wexlerae
Fusicatenibacter saccharivorans
Streptococcus thermophilus
Faecalibacterium prausnitzii
Taxa significantly (p <= 0.05) enriched in microbiomes from target disease as compared to non-diseased (GD) /other diseases (DS)
Taxa significantly (p <= 0.05) depleted in microbiomes from target disease as compared to non-diseased (GD) /other diseases (DS)
Taxa enriched in microbiomes from target disease as compared to non-diseased (GD) /other diseases (DS) with marginal significance ( 0.05 < p <= 0.1)
Taxa depleted in microbiomes from target disease as compared to non-diseased (GD) /other diseases (DS) with marginal significance ( 0.05 < p <= 0.1)
Significant (p<=0.05) difference between two signatures
Non-significant/absent among the top 20 taxa in that cluster

**Fig. S11. SHAP-based microbiome stratification reveals reproducible disease-specific signatures for CVD, IBS and SCZ. A-C**. Disease-specific microbiome signatures identified for cardiovascular diseases (CVD) (**A**), irritable bowel syndrome (IBS) (**B**) and Schizophrenia (SCZ) (**C**), based on clustering of microbiome SHAP profiles generated using the corresponding disease-specific MifRix ensemble models. For each disease, the left panel shows principal component analysis (PCA) of SHAP values from the testing cohort, with microbiome signatures defined by k-means clustering. Microbiomes from the independent validation cohort were projected onto the PCA space derived from the testing cohort to evaluate the reproducibility of the identified signatures. The upper middle panels compare predicted risk scores for ten diseases (Alzheimer's Disease (AD), Colorectal Cancer (CRC), CVD, inflammatory bowel disease (IBD), IBS, Multiple Sclerosis (MS), Parkinson's Disease (PD), SCZ, Type-II Diabetes (T2D) and Major Depressive Disorder (MDD)) across the identified microbiome signatures. Red dashed lines indicate the disease-specific risk threshold, and asterisks denote significant differences between signatures ($p \leq 0.05$). The lower middle panels show the proportion of testing and validation microbiomes assigned to each signature. Right panels summarize signature-associated microbial taxa that were enriched or depleted relative to microbiomes from the target disease versus non-diseased controls (GD) or other diseases (DS) in both testing and validation datasets. Heatmap cells are colored according to the direction and statistical significance of differential abundance: Dark magenta indicates significant enrichment (Hedges' $g > 0$ & $p \leq 0.05$); dark blue indicates significant depletion (Hedges' $g < 0$ & $p \leq 0.05$); light pink indicates marginal enrichment (Hedges' $g > 0$ & $0.05 < p \leq 0.1$); light blue indicates marginal depletion (Hedges' $g < 0$ & $0.05 < p \leq 0.1$); and white denotes non-significant or absent among the top 20 taxa for that signature.

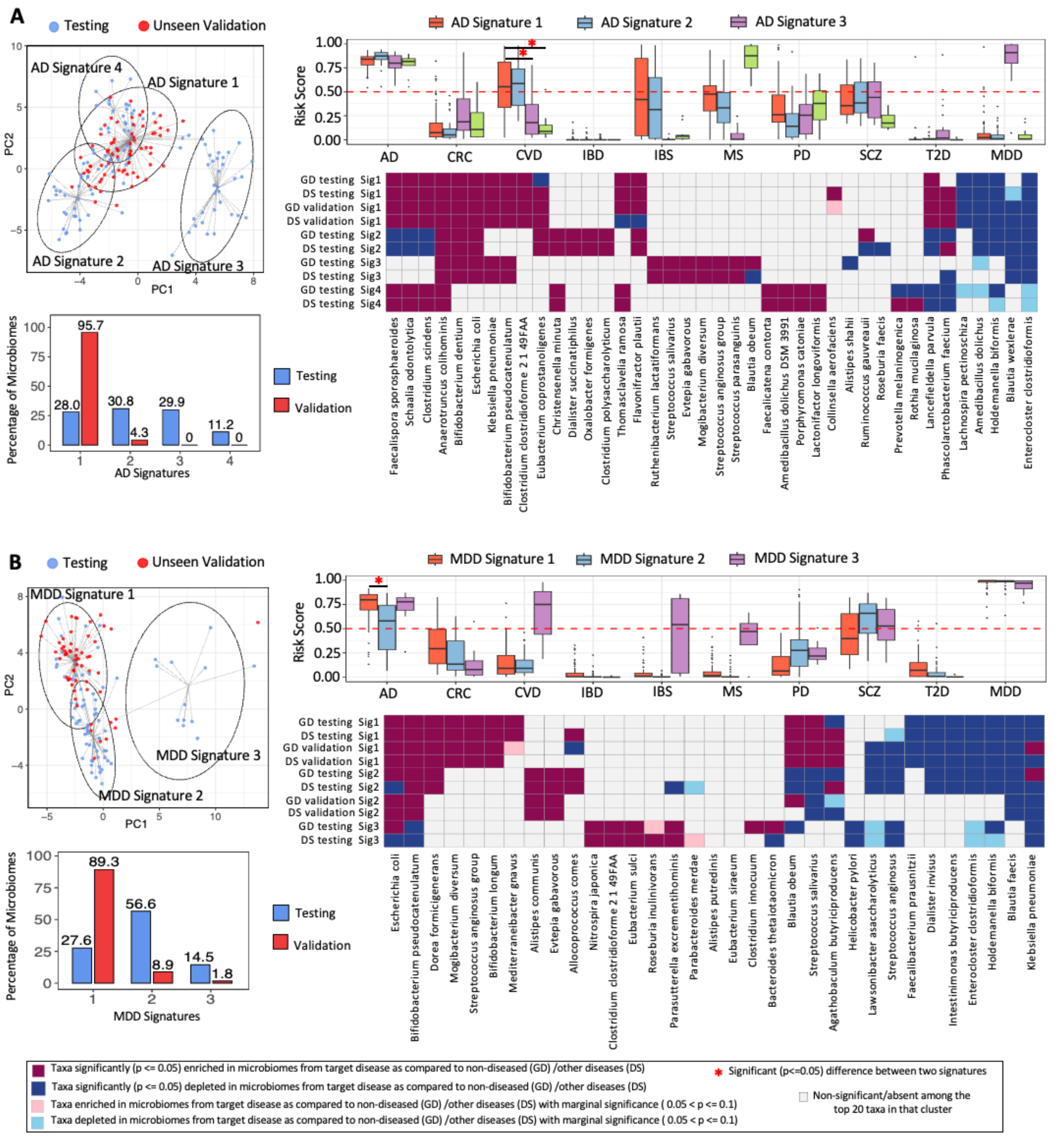


**Fig. S12. A-B.** Disease-specific microbiome signatures identified for Alzheimer's Disease (AD) (**A**) and Major Depressive Disorder (MDD) (**B**) based on clustering of microbiome SHAP profiles generated using the corresponding disease-specific MifRix ensemble models. For each disease, the left panel shows principal component analysis (PCA) of SHAP values from the testing cohort (in blue points), with microbiome signatures defined by k-means clustering. Microbiomes from the independent validation cohort (as red points) were projected onto the PCA space derived from the testing cohort to assess signature reproducibility. The upper middle panels compare predicted risk scores for ten diseases (AD, colorectal cancer (CRC), cardiovascular disease (CVD), inflammatory bowel disease (IBD), irritable bowel syndrome (IBS), MS, Parkinson's disease (PD), schizophrenia (SCZ), type-II diabetes (T2D) and major depressive disorder (MDD)) across the identified microbiome signatures. Red dashed lines indicate the disease-specific risk threshold, and asterisks denote significant differences between signatures ($p \leq 0.05$). The lower middle panels show the proportion of testing and validation microbiomes assigned to each signature. Right panels summarize signature-associated microbial taxa that were enriched or depleted relative to microbiomes from the target disease versus non-diseased controls (GD) or other diseases (DS) in both testing and validation datasets. Heatmap cells are colored according to the direction and statistical significance of differential abundance: dark magenta indicates significant enrichment (Hedges' $g > 0$ & $p \leq 0.05$); dark blue indicates significant depletion (Hedges' $g < 0$ & $p \leq 0.05$); light pink indicates marginal enrichment (Hedges' $g > 0$ & $0.05 < p \leq 0.1$); light blue indicates marginal depletion (Hedges' $g < 0$ & $0.05 < p \leq 0.1$); and white denotes non-significant or absent among the top 20 taxa for that signature.

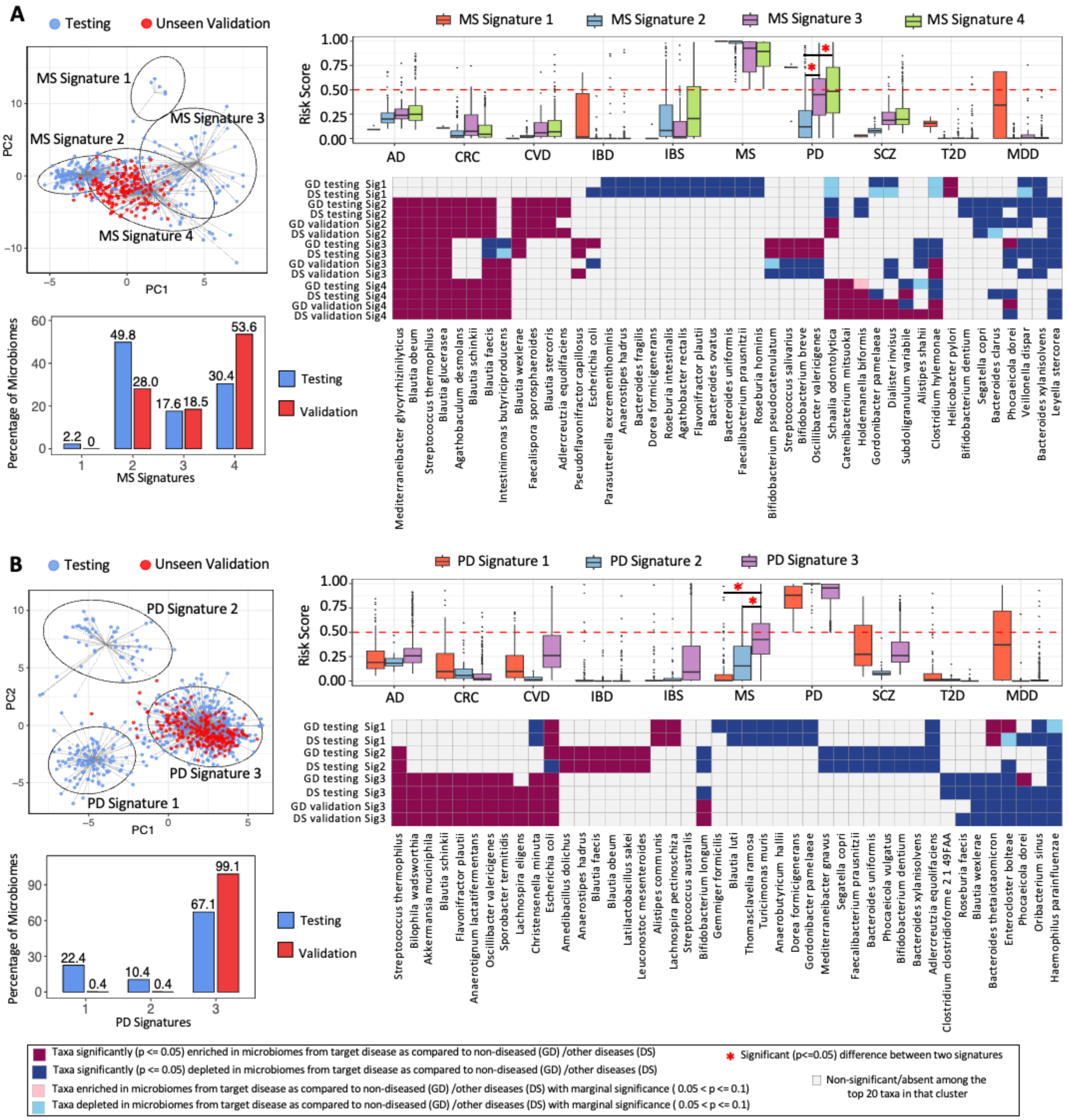


**Fig. S13. A-B.** Disease-specific microbiome signatures identified for Multiple Sclerosis (MS) (**A**) and Parkinson's Disease (PD) (**B**) based on clustering of microbiome SHAP profiles generated using the corresponding disease-specific MifRix ensemble models. For each disease, the left panel shows principal component analysis (PCA) of SHAP values from the testing cohort (in blue points), with microbiome signatures defined by k-means clustering. Microbiomes from the independent validation cohort (as red points) were projected onto the PCA space derived from the testing cohort to assess signature reproducibility. The upper middle panels compare predicted risk scores for ten diseases (AD, CRC, CVD, IBD, IBS, MS, PD, SCZ, T2D and MDD) across the identified microbiome signatures. Red dashed lines indicate the disease-specific risk threshold, and asterisks denote significant differences between signatures ($p \leq 0.05$). The lower middle panels show the proportion of testing and validation microbiomes assigned to each signature. Right panels summarize signature-associated microbial taxa that were enriched or depleted relative to microbiomes from the target disease versus non-diseased controls (GD) or other diseases (DS) in both testing and validation datasets. Heatmap cells are colored according to the direction and statistical significance of differential abundance: dark magenta indicates significant enrichment (Hedges' $g > 0$ & $p \leq 0.05$); dark blue indicates significant depletion (Hedges' $g < 0$ & $p \leq 0.05$); light pink indicates marginal enrichment (Hedges' $g > 0$ & $0.05 < p \leq 0.1$); light blue indicates marginal depletion (Hedges' $g < 0$ & $0.05 < p \leq 0.1$); and white denotes non-significant or absent among the top 20 taxa for that signature.

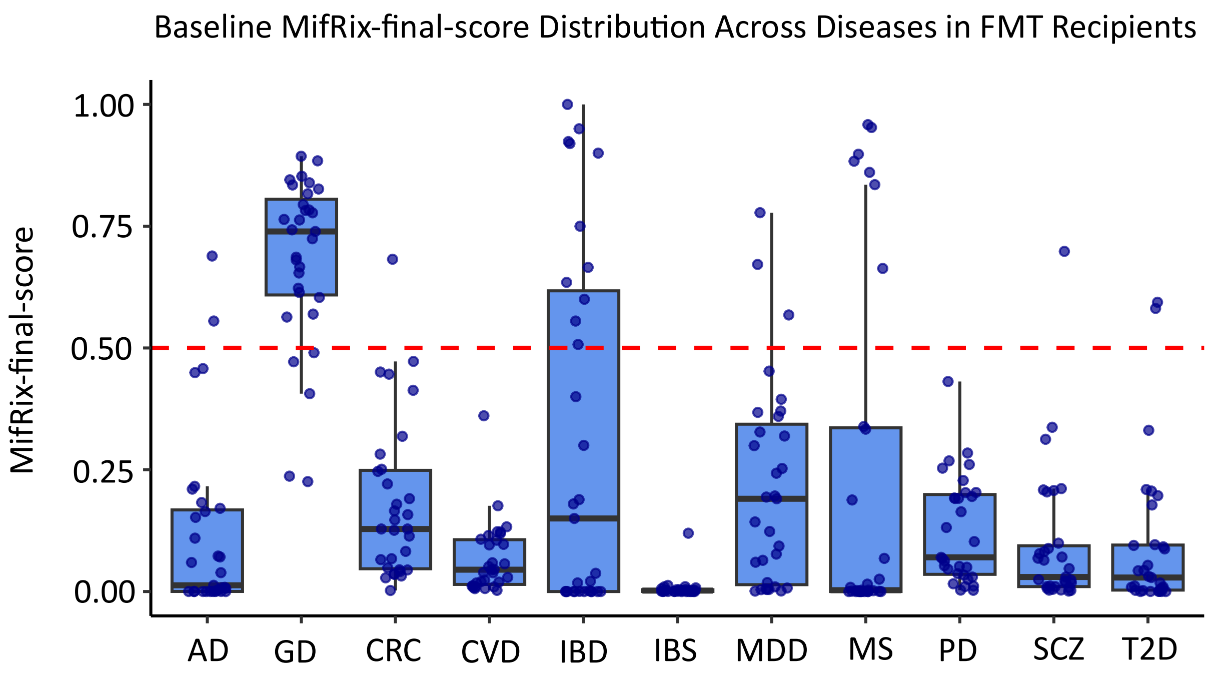


**Fig. S14. Generic Disease Signatures Best Captured the Baseline Disease State, Followed by IBD-Specific Models.** Baseline gut microbiome samples from 69 IBD patients enrolled across three independent FMT studies were analyzed using the MifRix framework to estimate generic disease risk as well as disease-specific risk-scores. Boxplots show the distribution of predicted risk-scores generated using 11 ensemble predictors, including Generic Disease (GD) risk, Alzheimer's disease (AD), colorectal cancer (CRC), cardiovascular disease (CVD), inflammatory bowel disease (IBD), irritable bowel syndrome (IBS), major depressive disorder (MDD), multiple sclerosis (MS), Parkinson's disease (PD), schizophrenia (SCZ), and type 2 diabetes (T2D). Each point represents an individual baseline microbiome sample. Boxes indicate the interquartile range (IQR), the central line represents the median, and whiskers extend to 1.5 × IQR. The red dashed line denotes the decision threshold (risk score = 0.5), above which samples are classified as high risk for the corresponding disease by MifRix. Among the evaluated diseases, baseline IBD samples exhibited the highest predicted risk for the Generic Disease (GD) model, followed by the IBD model, highlighting the predominance of shared dysbiosis-associated microbial signatures in the pre-FMT microbiome.
